# Multiscale harmonization, semantic integration, and immersive exploration of single-cell data in support of novel biological insights

**DOI:** 10.64898/2026.07.07.737090

**Authors:** Andreas Bueckle, Chenchen Zhu, Alex Y. H. Wong, Archibald Enninful, Yang Miao, Negin Farzad, Maria Pedersen, Courteney Mattison, Nicholas Sloan, Jason Mares, Cheng Xing, Bruce W. Herr, Juhi Khare, Yash Kumar, Keyur Parekh, Siddhi Chavan, Paean Luby, Ushma Patel, Yashvardhan Jain, John Hickey, Gary D. Bader, Hemali Phatnani, Vilas Menon, Rong Fan, Peter K. Sorger, Michael Snyder, Katy Börner

**Affiliations:** Department of Intelligent Systems Engineering, Indiana University, Bloomington, IN, USA; Department of Genetics, Stanford University, Stanford, CA, USA; Laboratory of Systems Pharmacology, Harvard Medical School, Boston, MA, USA; Ludwig Centre at Harvard, Harvard Medical School, Boston, MA, USA; School of Engineering & Applied Science, Yale University, New Haven, CT, USA; Department of Biomedical Engineering, Duke University, Durham, NC, USA; Center for Genomics of Neurodegenerative Disease, New York Genome Center, New York, NY, USA; Department of Neurology, Columbia University Irving Medical Center, New York, NY, USA; The Donnelly Centre, University of Toronto, Toronto, ON, Canada; Department of Molecular Genetics, University of Toronto, Toronto, ON, Canada; Princess Margaret Research Institute, University Health Network, Toronto, ON, Canada; Department of Computer Science, University of Toronto, Toronto, ON, Canada; Lunenfeld-Tanenbaum Research Institute, Toronto, ON, Canada; Canadian Institute for Advanced Research (CIFAR), Toronto, ON, Canada; Center for Translational and Computational Neuroimmunology, Department of Neurology, Columbia University Irving Medical Center, New York, NY, USA; Berlin Institute of Health at Charité, Universitätsmedizin Berlin, Berlin, Germany

## Abstract

Single-cell atlassing efforts like the Human BioMolecular Atlas Program (HuBMAP) and Cellular Senescence Network (SenNet) produce multiscale datasets across the healthy, adult, human body but are typically explored on 2D screens with limited 3D affordances, even though understanding cell location within tissue, organ, and body requires spatial reasoning across many orders of magnitude. The Human Reference Atlas (HRA) provides standard terminologies and a Common Coordinate Framework (CCF) for harmonizing this data spatially and semantically. Building on the **HRA Organ Gallery in virtual reality (VR)**, we present **“HRA: Powers of Ten**,**”** which integrates, harmonizes, and visualizes biological data from these efforts in VR using a **Multiscale Elevator System** that lets users descend, like riding an elevator through an inverted skyscraper, from a whole body view of 81 reference organs to datasets across 5 organs (lymph node, brain, large intestine, small intestine, liver), 5 assay types (CODEX, Visium, v-CyCIF, Xenium, SBF-SEM), and 4 spatial scales. Shared by Data Providers, these VR scenes enable biological insights into senescence patterns, cellular neighborhoods, 3D tissue reconstruction, and subcellular liver architecture. A standard operating procedure supports adding further datasets. Freely available on the Meta Store to over 20 million headsets, the application, data, code are open-source.

## Introduction

Single-cell atlassing portals such as HuBMAP^1,2^ and SenNet^3^ publish datasets across scales for multiple organs of the healthy adult human body for both sexes, multiple ethnicities, and across adult ages. The Canadian Institute for Advanced Research (CIFAR) MacMillan Multiscale Human (cifar.ca/research-programs/cifar-macmillan-multiscale-human) effort aims to build an explorable model of the human body across scales to drive advances in medicine and therapeutics. The data from these efforts can be interrogated semantically and spatially but only on 2D screens with limited 3D affordances^4^. Imagine, for example, examining the 3D location of a cell in the context of a tissue, inside of an organ, and as part of a human body, across multiple orders of magnitude. To be tackled effectively, 3D multiscale data and inquiries require 3D user interfaces (UIs).

In support of multiscale harmonization, the HRA^5^ provides terminologies and data structures for specimens, biological structures, and spatial positions of experimental datasets inside 3D reference organs of the adult human body^6^. Building on it, prior work has showcased the HRA Organ Gallery^7^, a VR application that lets users explore the HRA in immersive 3D. Here, we introduce “**HRA: Powers of Ten**,” a data integration and visualization application that uses the HRA Common Coordinate Framework (CCF)^6^ to harmonize, visualize, and explore data across 5 organs (lymph node, brain, large/small intestine, liver), 5 assay types (CO-Detection of IndEXing [CODEX]/PhenoCycler^8^, Visium [www.10xgenomics.com/platforms/visium], v-CyCIF^9^, Xenium^10^, and serial block-face scanning electron microscopy [SBF-SEM]^11^), and 4 scales (meters, centimeters, millimeters, microns). It features a **Multiscale Elevator System** where users can traverse the human body like an inverted skyscraper, with the Whole Body level at the top and each subsequent elevator floor shrinking the user by 1 order of magnitude, see details in the **Results > Multiscale Elevator System** section below. A total of 8 scenes are presented in the application; each contains data from a different Data Provider, all co-authors on this paper. The scenes are named after the effort from where the data originates (e.g., “hubmap”), the organ (“large_intestine”), the name(s) of the Data Provider(s) (“wong”), the prevalent metric scale (“100_microns”), and the same metric scale expressed as a power of 10, i.e., its Multiscale Elevator System floor (“10_4”). The **Results > Insights from VR** section describes biological insights obtained by Data Providers. The process of adding new datasets to the HRA Organ Gallery is described in the **Results > Adding new datasets** section and in a standard operating procedure (SOP)^12^.To enable readers without VR headsets to understand what data is presented in each scene, videos featuring voice-overs by the Data Providers and other relevant metadata are available at cns-iu.github.io/hra-organ-gallery-supporting-information/#videos. **Fig. 1** provides a high-level gallery view of all 8 scenes. **Fig. 2** provides detailed screenshots with illustrations of users.

**Fig. 1.**
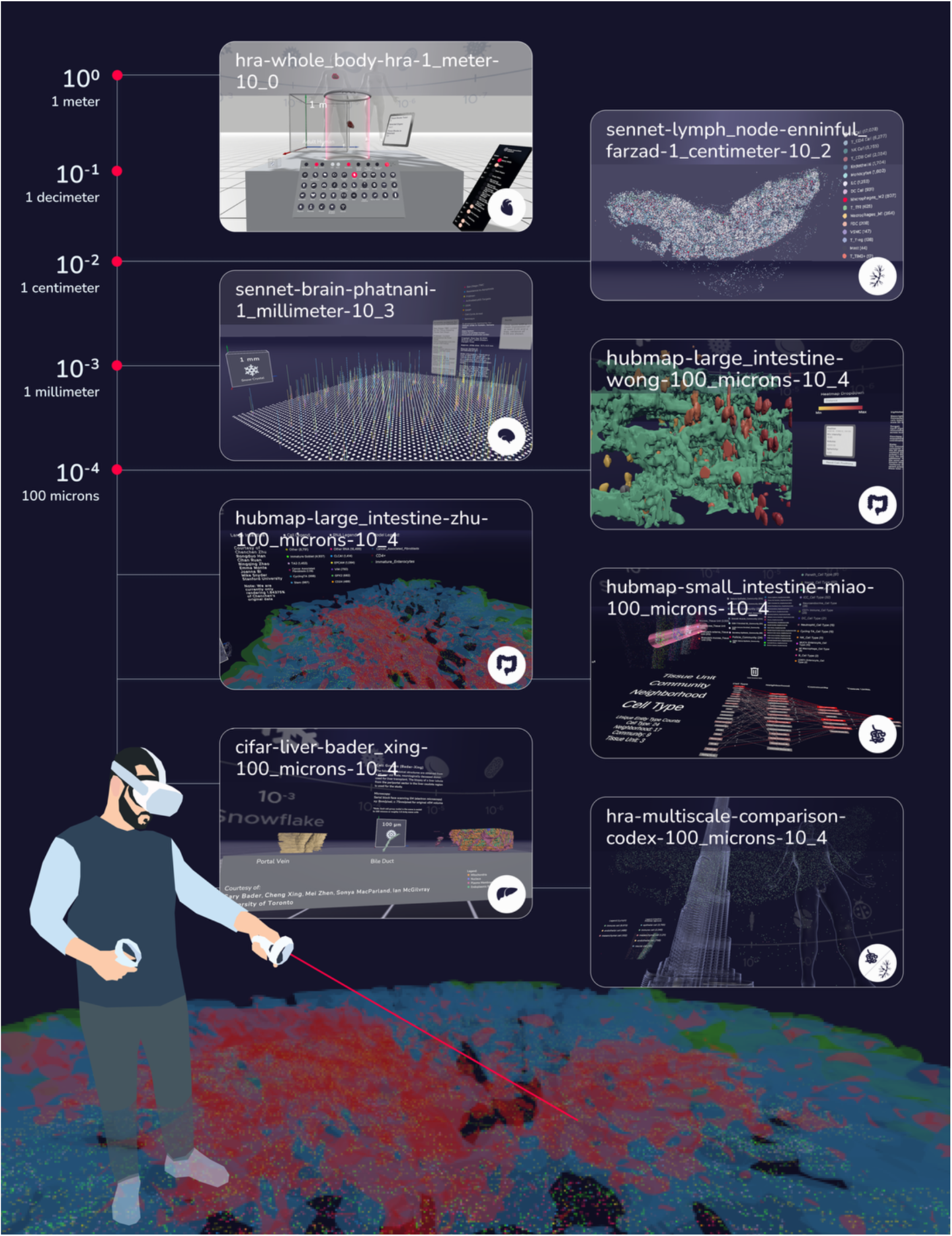
Overview of all scenes in HRA: Powers of Ten. VR enables Data Providers to perform HRA-based data harmonization and visualization across scales. Shown are screenshots from 8 scenes featured in this paper; these screenshots are connected to the corresponding level on the Multiscale Elevator System (on the left). At the bottom, a user is exploring a from the reconstructed 3D tissue map in the large intestine from **hubmap-large_intestine-zhu-100_microns-10_4**.

**Fig. 2.**
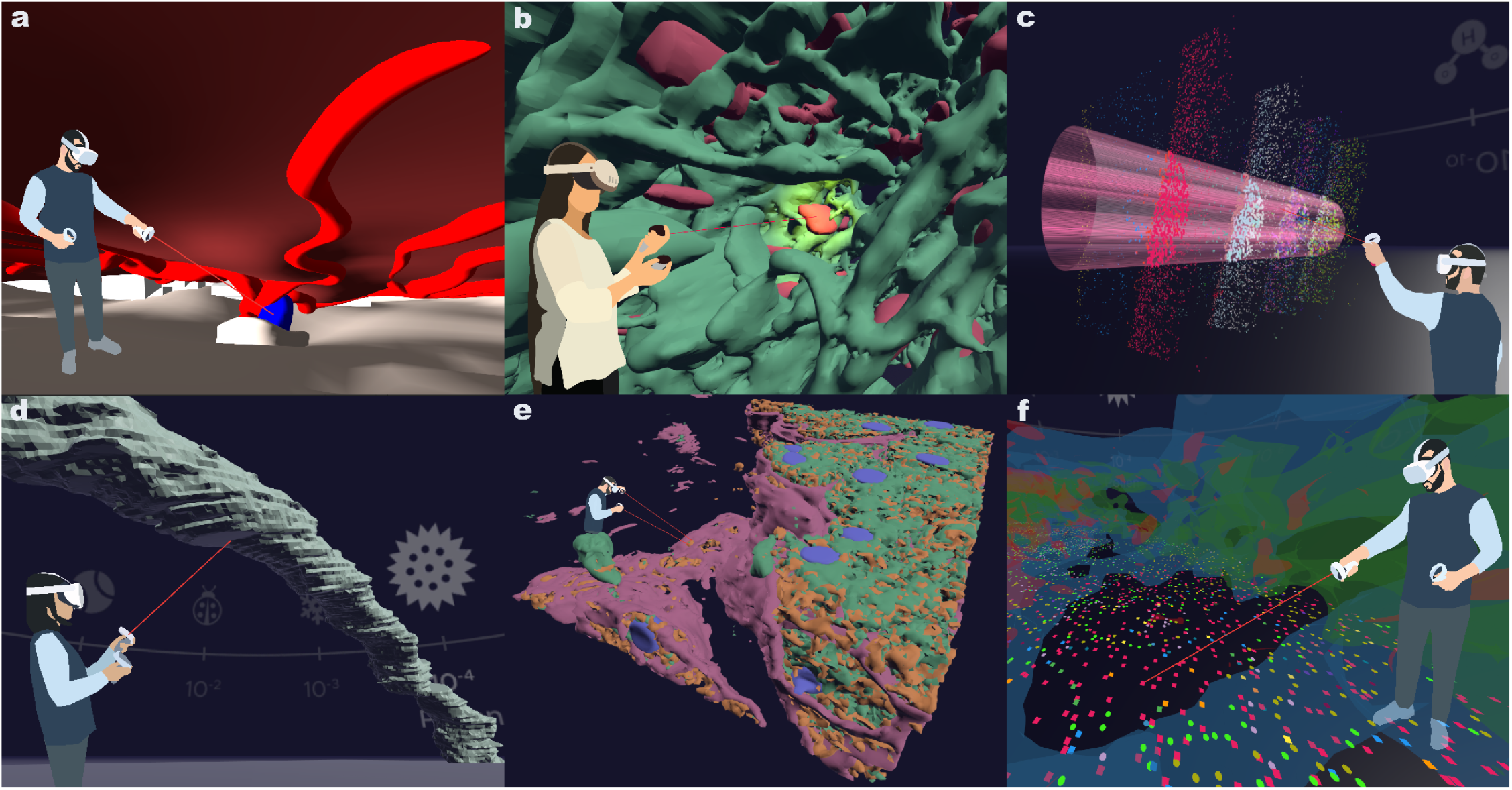
A tour of HRA: Powers of Ten. **a**, The maternal interface of the placenta with the umbilical cord being formed by blood vasculature in **hra-whole_body-hra-1_meter-10_0** after the user has scaled up the organ. **b**, The user inspects the CD8 cells with ID 72 in the cropped volume (light-sheet/v-CyCIF) in **hubmap-large_intestine-wong-100_microns-10_4. c**, The user places a search cylinder over the 4 aggregation layers in **hubmap-small_intestine-miao-100_microns-10_4. d**, The user points at the bile duct, enlarged 20x, in **cifar-liver-bader_xing-100_microns-10_4**. e, In the same scene, the user explores a 3D SBF-SEM volume with cell nuclei (blue), plasma membrane (pink), endoplasmic reticulum (green), and mitochondria (orange). **f**, The user inspects cell nuclei and RNA molecules on top of a 3D reconstruction of Xenium data from a polyp in the large intestine in **hubmap-large_intestine-zhu-100_microns-10_4**.

Implementation details with additional figures are provided in the **Methods** section. As part of the HRA Organ Gallery in VR, HRA: Powers of Ten is available for free through the Meta Store at www.meta.com/en-gb/experiences/hra-organ-gallery/5696814507101529 to over 20 million users owning affordable, portable, off-the-shelf Meta Quest headsets.

The development of 5 of the 8 scenes in this application was conducted collaboratively by 7 early-stage investigators (ESIs) and partially funded by a co-author Bueckle’s HuBMAP JumpStart Fellowship (hubmapconsortium.org/jumpstart-program/#andreas2024), awarded by the National Institutes of Health (NIH). All 7 ESIs are co-authors on this paper (Bueckle, Zhu, Wong, Enninful, Miao, Farzad, Xing); all served as Data Providers (scene names have ESI names in them); co-author Bueckle implemented all with his team as the Organ Gallery Lead (development lead). This work was conceived of during a JumpStart-funded workshop, see **Supplementary Note 1** and cns-iu.github.io/workshops/2024-10-24-jumpstart-workshop. 5 of the ESI authors attended this workshop.

### HRA: Powers of Ten

addresses an important gap in biomedical visualization. While commercial applications for viewing the human body are widely used to teach anatomy, none support multiscale data exploration from the whole body to (sub-)cellular resolution. All of these scales are critical to understanding human physiology. 3D stereoscopy, supported by increasingly high-resolution displays in VR headsets, enables accurate perception of depth and distances between different organs, cells, and subcellular entities (~2k pixels per eye for the Meta Quest 3 used in the HRA Organ Gallery); these also allow detailed views of spatial arrangements and size comparisons.

Further, natural input via VR controllers supports advanced interactions with data beyond traditional mouse and keyboard paradigms, e.g., grabbing, moving, rotating, and scaling cells and tissue regions with the user’s own hands. These effects are further enhanced by 2 features of the HRA Organ Gallery: 3D scale boxes (instead of 2D scale bars) with easy-to-recognize scale-specific objects as landmarks and virtual Elevator Transition Scenes to communicate scale changes (see the **Methods > Shared features between scenes** section for details). Where multiscale visualization has been implemented, it’s been so within a limited scope.

The vast majority of these applications are focused on genomics and allow users to compare genomes of varying size and phylogenies^13,14^. Some genome-oriented multiscale applications are more visualization-focused, such as the Multiscale Unfolding application that takes users from a genome’s chromosomes to its double helices and base pairs^15^. Multiscale visualizations are appreciably less common at the organ-level, and where they do exist they are frequently limited in their scope. For example, one VR prototype allows users to view a 3D mouse prostrate at the organ and sub-organ levels, with an additional quantitative histology level^16^. Another leveraged computed tomography to allow users to move from the whole organ scale down to the ion channel level with select muscles and bones^17^.

Prior work has also explored a variety of approaches for communicating and navigating scale, both within biological datasets and in broader educational and visualization contexts. For example, some have advocated for a Google Maps–like navigation paradigm where higher-resolution datasets are embedded within larger anatomical contexts and allow users to progressively zoom into regions of interest while maintaining spatial awareness across scales^17,18^. 2D experiences like the Nikon Universcale (www.nikon.com/company/corporate/sp/universcale/?ref=1), Kees Boeke’s influential illustrations in “Cosmic View: The Universe in Forty Jumps”^19^, and the “Powers of Ten” educational movie^20^ let users view objects across scales, but these are limited to non-stereoscopic web applications, static pages in a book, and non-interactive video.

## Results

### Multiscale Elevator System

To explore the human body, 10 orders of magnitude must be traversed in 3D, from the whole body down to proteins. HRA: Powers of Ten makes these accessible via a **Multiscale Elevator System** where the body is envisioned as an inverted skyscraper with an elevator taking the user from the Whole Body on the top floor down. Each level constitutes an order of magnitude scale change. Each dataset is visualized in its own *scene*, i.e., a closed 3D immersive space that the user can navigate, which is situated on a *level*, i.e., a discrete zoom magnification. Each level gets its “floor number,” or **power**, i.e., −4 in 10^−4^, from the exponent to base 10 of the scale of the dataset(s) featured. For example, in level 0, the Whole Body level, 1 meter in physical space corresponds to 10^0^ (=1) meters in virtual space, i.e., no magnification is applied. Likewise, in level −4, the Cell Groups level, 1 meter in physical space corresponds to 10^−4^ meters (=100 microns) in virtual space, which means a magnification of 4 orders of magnitude. Because this is a VR application, the user always has their own body as a reference, e.g., their extended arms. A level can have multiple scenes; for example, the 10^−4^ level has 5 scenes with data from 4 organs (lymph node, small intestine, large intestine, liver). To access them, a virtual **Elevator Panel** is featured in every scene that allows the user to move on to a new scene (see **Methods** section and **Fig. 3.d**). To further communicate scale differences between scenes, each level features an Icon of a real-world object, which is visible in the Skybox, i.e., the cube map texture that gives the illusion of a spherical sky, see **Methods** section and **Supplementary Fig. 1**. When switching to a scene on a different level, an optional Elevator Transition Scene is played where the user is placed inside a virtual elevator box and either sees the entire human body or, in lower scales, a collection of red blood cells as reference models given the appropriate scale difference (see **Supplementary Fig. 2** and the last video listed under cns-iu.github.io/hra-organ-gallery-supporting-information/#videos).

**Fig. 3.**
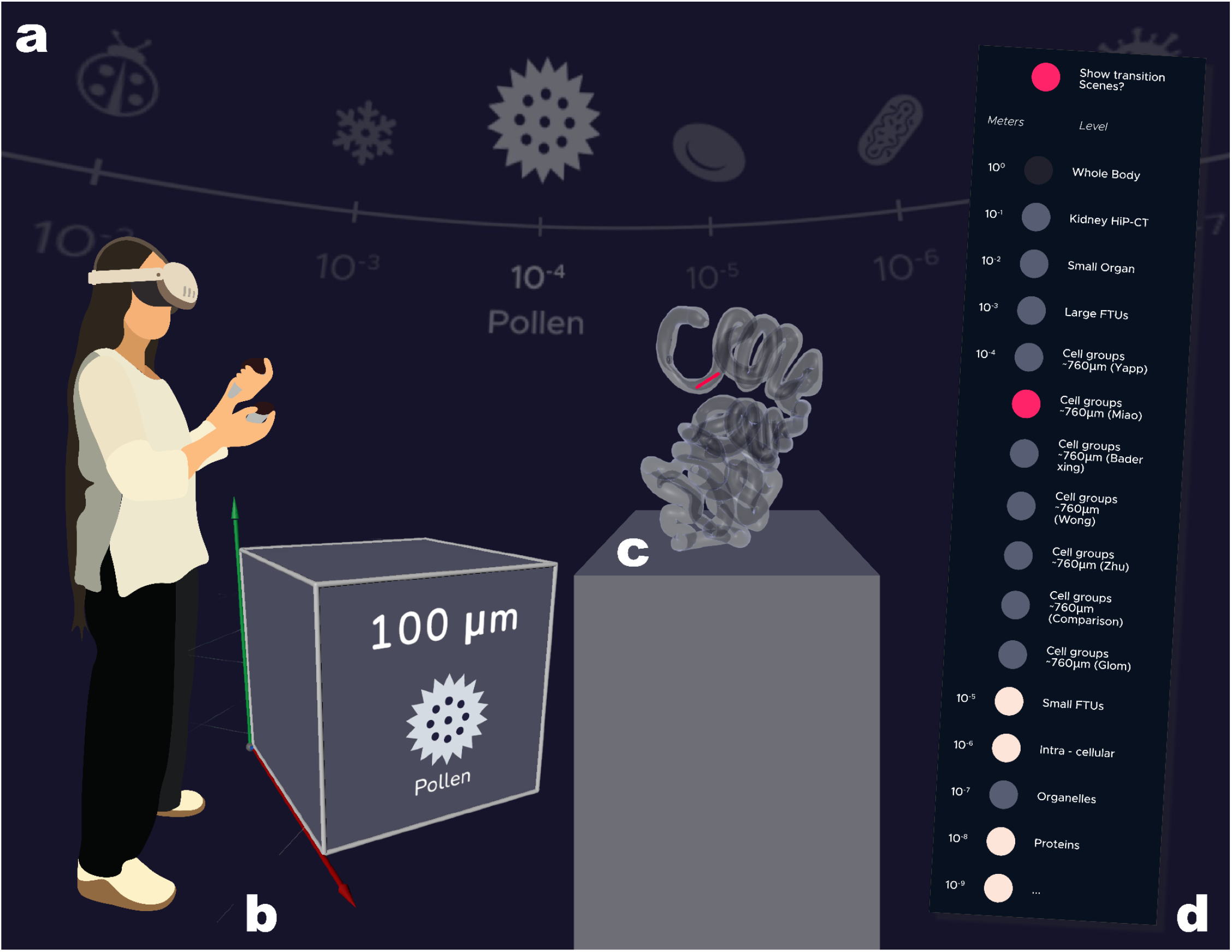
Shared features between scenes. **a**, Skyboxes with Icons of real-world objects. **b**, A Scale Cube with the Icon and metric scale of the scene (pollen, 10^−4^, 100 microns), and the name of the Icon spelled out. **c**, An Extraction Site Component for a tissue block (sample: HBM787.FHZM.948, dataset: HBM573.SWGH.988) in the small intestine from **hubmap-small_intestine-miao-100_microns-10_4. d**, An Elevator Panel allowing users to travel to scenes across the human body.

### Insights from VR

The 8 scenes presented in this paper are spread across 4 levels: the Whole Body level with 81 3D reference organs of the HRA v2.5 (10^0^, 1 meter), Small Organs (10^−2^, 1 centimeter), Large FTUs (10^−3^, 1 millimeter), and Cell Groups (10^−4^, 100 microns). Each scene gets its name from (a) the funded effort that the data originates from, (b) the organ, (c) the last name of the Data Provider(s), and (d) the level at which the data was integrated.

**Fig. 4** presents details on the datasets used in HRA: Powers of Ten scenes. Implementation details for all scenes are provided in the **Methods** section.

**Fig. 4.**
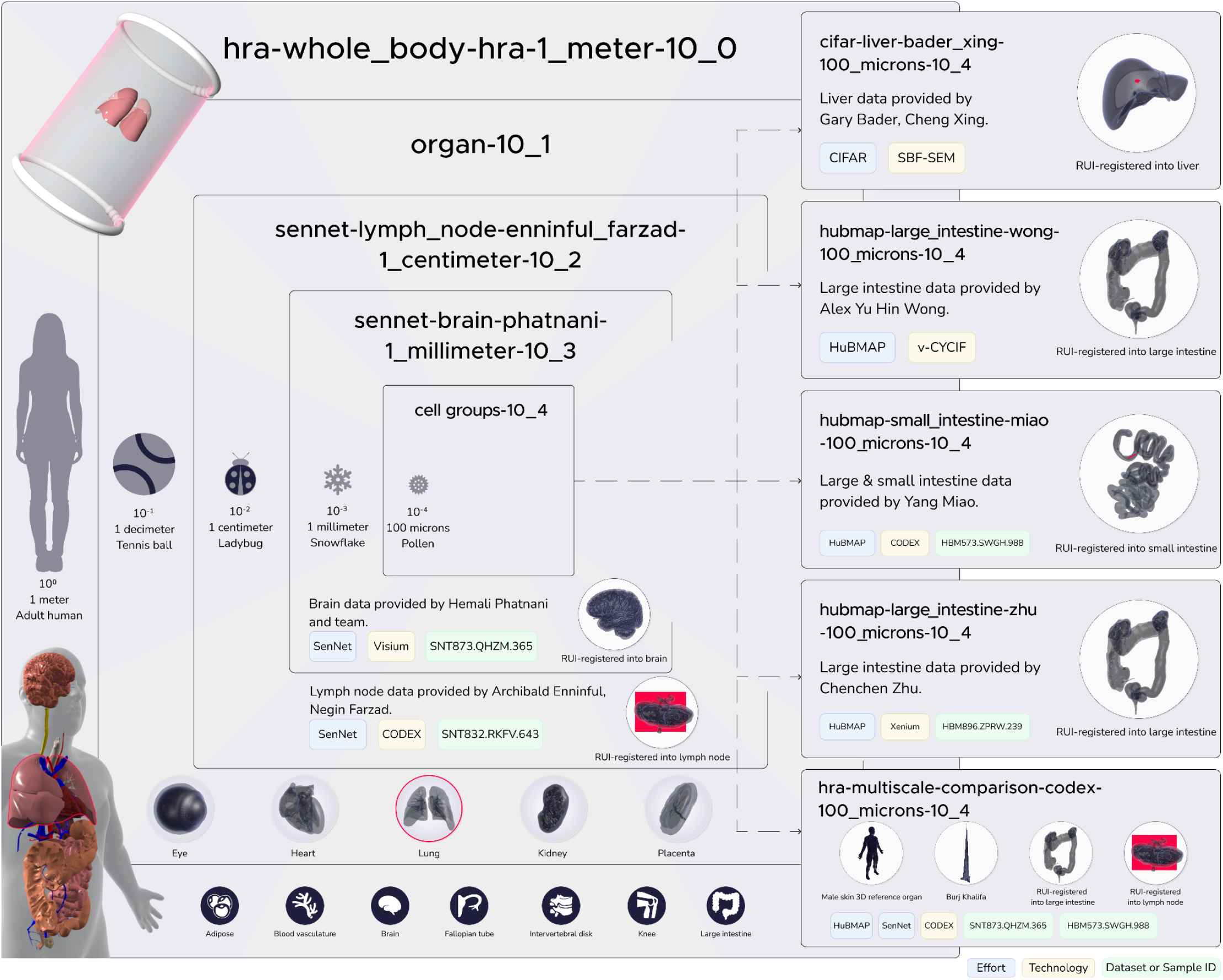
Metadata for the 8 scenes in this paper in the context of the Multiscale Elevator System of HRA: Powers of Ten.

### hra-whole_body-hra-1_meter-10_0

*(a whole body introduction to the HRA)*: As of September 2026, the start scene of the HRA Organ Gallery contains 81 3D reference organs of HRA v2.5 alongside 2,762 tissue blocks representing 13,344 datasets from 1,021 donors, registered by 54 tissue data providers with the Registration User Interface (RUI)^4^. Using a virtual keyboard with Icons, the user can select an organ (plus sex and laterality if available) to load a high-resolution version of a 3D organ to move, rotate, and scale while inspecting the tissue blocks in their native (or relative) size. They can also inspect anatomical details in the organ (such as the umbilical cord shown in **Fig. 2.a**). A detailed view is available in **Supplementary Fig. 3**.

### sennet-lymph_node-enninful_farzad-1_centimeter-10_2

*(spatial patterns of senescent cells)*: Interactions between senescent^3^ cells in human lymph nodes are easy to miss in 2D. HRA: Powers of Ten features a CODEX dataset^21^ obtained from a 86-year old male donor with SenNet ID SNT832.RKFV.643 featuring 36,964 cells of 16 cell types shown (see **Supplementary Fig. 4**). The scene illustrates that lymph nodes are highly structured tissues with distinct cellular regions, including B-cell-rich follicles and T-cell-dominant interfollicular zones. Additional immune cells, such as macrophages, monocytes, and dendritic cells are distributed in spatial patterns that reflect their roles in immune surveillance and response. This enables a more intuitive 3D exploration by making their spatial patterns and interactions with other cells within their microenvironment explicit. Further, this immersive approach makes it easier to examine the relationship between immune cells and vasculature as well as stromal cells in the lymph node. A brush and brush-and-link function to highlight all occurrences of a cell type across the tissue, the user can map individual immune cells and their spatial relationships within the tissue. Distinct anatomical regions are visible, including B cell-rich follicles and T cell-dominated interfollicular zones, along with macrophages, dendritic cells, and vascular cells. Looking ahead, VR holds promise for deeper understanding of complex tissue microenvironments particularly within the context of cancers. With spatial proteomic data generated from serial tissue sections, there is hope to construct full 3D atlases of lymph nodes to study lymphomas (specifically Diffuse Large B Cell Lymphoma ^22^ and Angioimmunoblastic T Cell Lymphoma^23^) in unprecedented detail via HTAN (see humantumoratlas.org/center/hta209). VR can help visualize tumor cells and track their evolution within the tumor microenvironment. A detailed view is available in **Supplementary Fig. 4**.

### sennet-brain-phatnani-1_millimeter-10_3

(*simultaneous comparison of 8 senescence hallmarks in 3D space)*: The scene features a Visium slide of dorsolateral prefrontal cortex tissue from the brain of a 74-year-old female donor (dataset ID SNT873.QHZM.365). It contains a 3D spike plot with color-coded senescence hallmark expressions for senescent cells. The capture area of the Visium slide is 6.5 × 6.5 millimeters, magnified by 3 orders of magnitude to 6.5 x 6.5 meters. There are a total of 4,992 spots per capture area; each spot is 55 microns in diameter with a 100 microns center-to-center distance between spots. Each spot represents gene activity from small cell groups, visualized with colored lines indicating levels of aging-related gene signatures. Different colors correspond to curated gene sets representing 8 senescence hallmarks: Resistance.to.Apoptosis, DDR, SASP, Cell.Cycle.Arrest (see related publication^24^ for details on these four); Fridman^25^; Activated.p.Targets^26^, Senmayo^27^; and those curated by the San DiegoTissue Mapping Center. Lines above or below the plane of a cell spot show enrichment or depletion of these genes. This visualization helps reveal spatial patterns and relationships in senescence.

Identifying patterns, trends, and outliers in these 39,936 spatially explicit expression markers is non-trivial. Using data visualizations such as the 3D spike plot presented here, users can investigate spatial patterns that may indicate subclinical senescence, emerging senotypes, or localized senescence microenvironments (SMEs). Do zones of elevated SASP border regions with early DDR or cell-cycle arrest? Do these arrangements indicate tissue-specific aging pathways? Through immersive, multi-hallmark research, this scene enables hypothesis generation and the development of computational models that capture spatial relationships not easily detectable in 2D. Additional 2D plots enable auxiliary data visualization with scatter graphs with individual senescence hallmark values. A detailed view is available in **Fig. 5**.

**Fig. 5.**
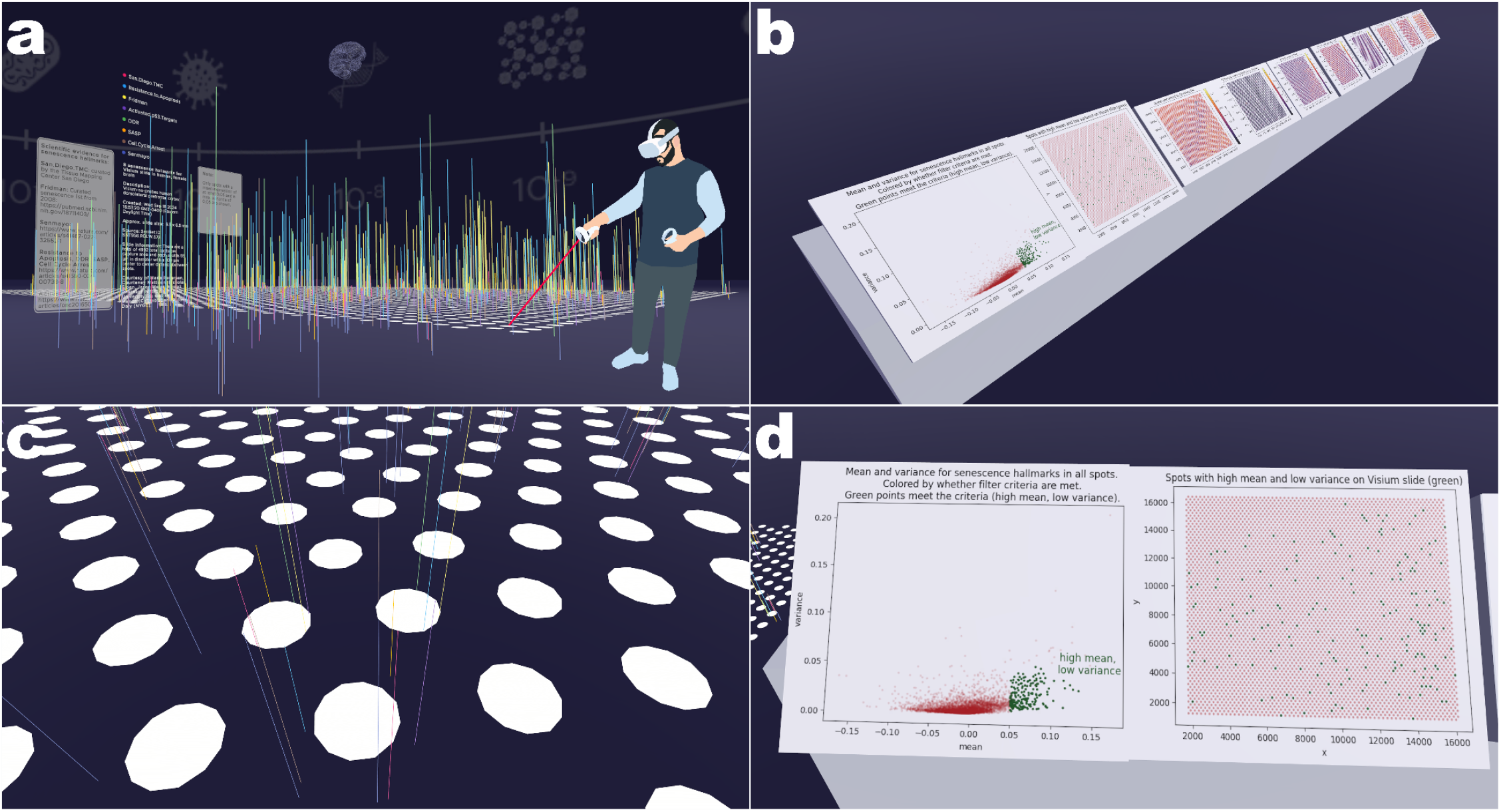
Visualization of cellular senescence hallmarks in SenNet brain tissue in sennet-brain-phatnani-1_millimeter-10_3. **a**, 3D spike plot for 8 senescence hallmarks (color-coded) above or below the plane of the cell spots. **b**, 2D plot gallery with 10 plots. **c**, Detail view of several cell spots, some of them with senescence hallmark expression lines below the plane of the spot. **d**, Detail view of 2 auxiliary graphs (left: mean [x] and variance [y] for senescence hallmark expressions in each cell spot; cell spots in green, i.e., high mean and low variance, get lines for their associated senescence hallmarks; right: the same data but with each cell spots x and y-position in scatter graph).

### hubmap-large_intestine-wong-100_microns-10_4

*(Volumetric cyclic immunofluorescence (v-CyCIF)* ^9^ *reveals 3D topology of cells and surrounding nerves in clinical specimens)*: How can multiscale data with many different formats (tabular, nested/hierarchical, 2D, 3D) be integrated into a coherent exploration and analysis space^28^? How can this inherently static data “come alive,” become more interactive, and multi-modal for the user? To explore these questions, this scene features 2 complementary volumetric datasets from the human large intestine. The first is a light-sheet^29^ microscopy image crop featuring ~200 CD8^+^ conventional T cells (Tcon) and βIII-tubulin expressing (TUBB3) enteric nerves. The second is a high resolution confocal dataset ^30^ with blood vessels (CD31, endothelial cells), epithelial cells (Claudin-7), in addition to T cells and nerves. In the light-sheet dataset, CD8^+^ Tcon cells and nerve structures are mapped in 3D, which shows most immune cells in direct physical contact with nerves, while some migrate towards the lumen. Users can interactively inspect cells by grabbing, moving, and rotating them in 3D using their VR controllers; they can visualize properties (distance to nearest nerve ending, volume, minimum intensity, and sphericity) via a heatmap. As a result, HRA: Powers of Ten uses VR visualization to transform static image volumes into immersive, manipulable representations of tissue organization. Together, this scene combines the imaging depth of light-sheet microscopy with the spatial resolution of confocal microscopy, which enables multiscale analysis of intact tissue architecture. The approach uses v-CyCIF^*9*^ to integrate multiple markers, which enables detailed spatial analysis of immune, epithelial, and vascular structures. In 2D, the 3D morphology of individual cells remains obscured by their proximity within dense cellular neighborhoods. VR supports an “exploded view” that greatly enhances inspection capabilities. Users can virtually separate and examine 3D T cells in detail and discard cells with poor segmentation quality. A detailed view is available in **Fig. 6** and **Fig. 2.b**.

**Fig. 6.**
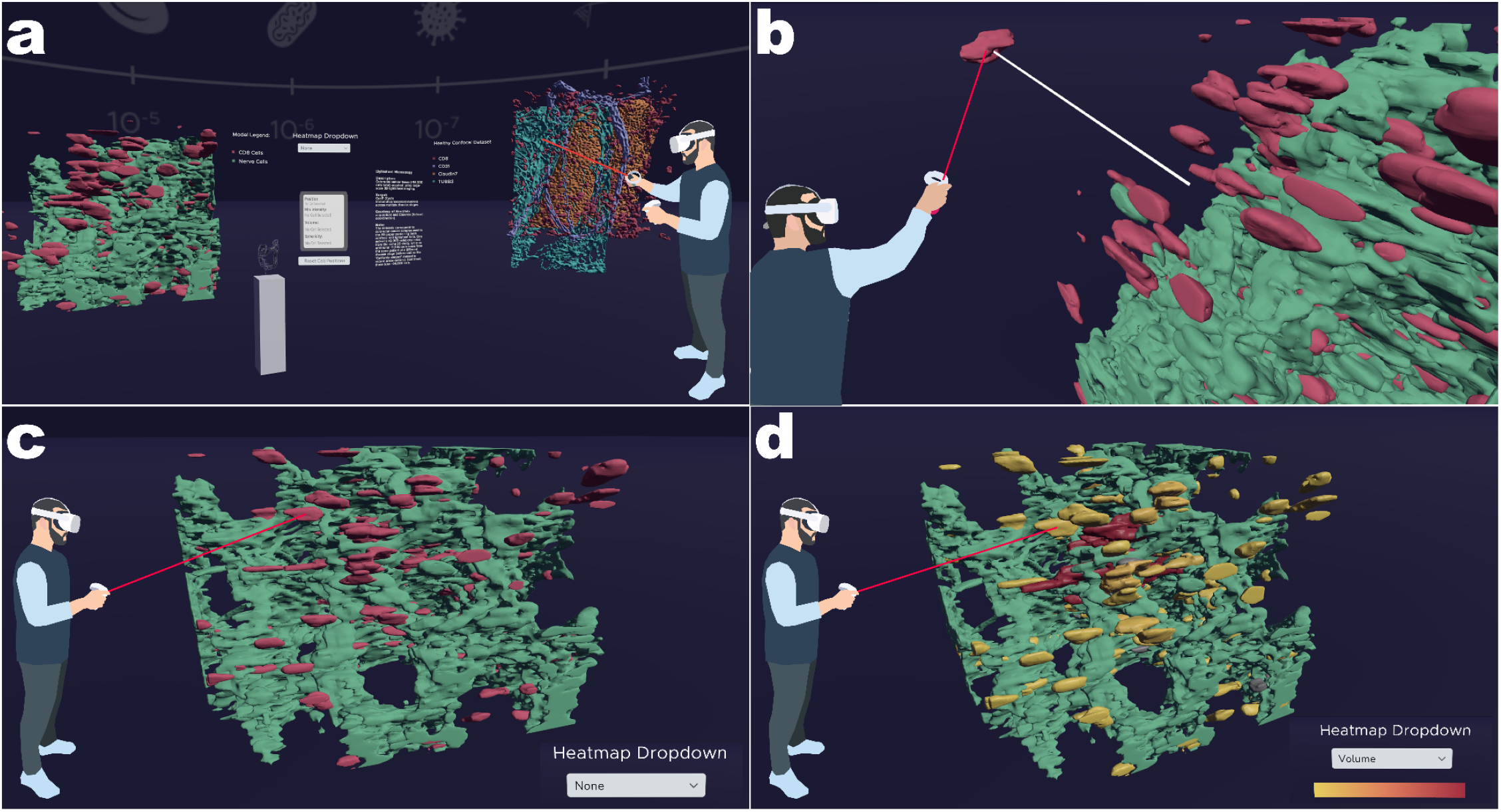
a, Overview of light-sheet and confocal microscopy datasets in shared 3D space in hubmap-large_intestine-wong-100_microns-10_4. **b**, The user grabs a cell; a visual tether line (white) is visible. **c**, detail shot of the light-sheet model with dropdown = None. **d**, Same detail shot with dropdown = volume. The color scheme used here is *YlOrRd* from Color Brewer (colorbrewer2.org/#type=sequential&scheme=YlOrRd&n=3).

### hubmap-small_intestine-miao-100_microns-10_4

*(layered presentations of cellular neighborhoods)*: The complexity of multi-scale cellular neighborhoods and tissue organization^31,32^ in the human body is difficult to appreciate, understand, and interpret in static views. HRA: Powers of Ten enables researchers to visualize and overlay cellular neighborhoods across multiple levels of aggregation, i.e., from the local microenvironment to entire tissue units. This scene features a VR visualization of a representative small-intestine tissue region^31^ acquired using CODEX (dataset ID HBM573.SWGH.988). The tissue is displayed across 4 ascending levels of aggregation: cell type, neighborhood, community, and tissue unit, thus, the same cell is represented across 4 interconnected levels of aggregation. At the cell-type level, individual cells are shown by their annotated identities. At higher levels, cells are grouped based on the composition of their surrounding neighbors, with each scale using a different neighborhood range to reveal tissue organization at increasing spatial breadth. At the neighborhood scale, compartmentalized local microenvironments that support specific tissue functions are captured. The broadest scale, tissue unit, aligns with known intestinal histology, spanning structures from the mucosa to the muscularis externa. The visualization of different scales allows users to explore finer details, e.g., identifying CD4^+^ and CD8^+^ T cells within unique multicellular environments. An alluvial diagram further enables simultaneous multi-scale analysis and quantitative network visualization by linking cellular distributions across levels. Additionally, an interactive selection cone that highlights the same cell across different aggregation levels further enhances spatial interpretation and enables users to connect local cellular identity with higher-order tissue architecture. This scene features 3,252 cells at 4 aggregation levels. A detailed view is available in **Fig. 7** and **Fig. 2.c**.

**Fig. 7.**
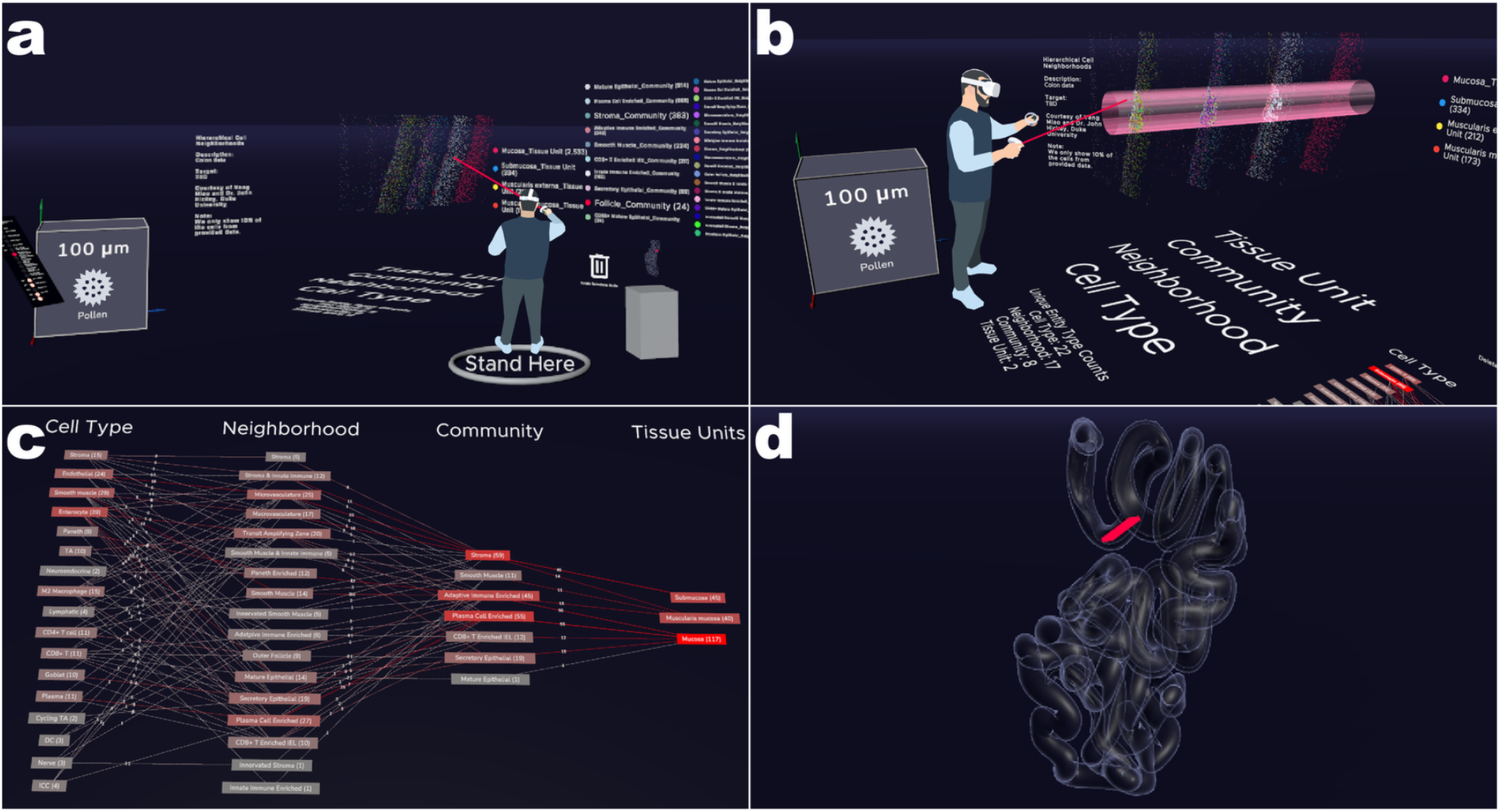
Cellular neighborhood visualization VR in hubmap-small_intestine-miao-100_microns-10_4. **a**, A “Stand Here” sign that directs the user to a good starting point with a view of all features in the scene. **b**, 4 layers of aggregation with a selection cylinder and a text field that shows the number of unique cell types, neighborhoods, communities, and tissue units (similar to what is shown in **Fig. 2.c**). **c**, Overhead screenshot of the alluvial diagram with the number of cells per types, neighborhoods, communities, and tissue units. **d**, Extraction Site Component showing the spatial origin of dataset HBM573.SWGH.988.

### hubmap-large_intestine-zhu-100_microns-10_4

*(integrating 3D tissue, 2D cells, and RNA molecules into a coherent, multiscale zoom)*: Due to their flatness, 2D visualizations can only capture so much detail about the stereotyped organization of tissue. By creating an immersive environment that enables visualization across multiple biological scales (tissues, cells, and molecules), VR helps users make more informed identifications of cell types and better understand putative crosstalks between different cell types, e.g., the co-localization of signaling molecules and ligand-receptor pairs, particularly among diverse tissue-resident immune cells. This enables scientists to better assess and refine their 3D tissue reconstruction models. This HRA: Powers of Ten scene features a 3D model of a human pre-cancerous large intestine polyp (sample ID HBM896.ZPRW.239). The scene integrates 3 data types: (1) a 3D reconstruction of the polyp from 29 serial thin sections were assayed using 10x Xenium^10^ assay, a highly multiplexed spatial method for identifying RNA molecules *in situ*; the reconstruction was achieved with Space-map^33^, a method optimized for reconstructing 3D maps from serial tissue sections, to generate the anatomically faithful 3D model presented here; (2) the same 29 layers as lists of 2D cells with x-position, y-position, and cell type labels; and (3) RNA molecules as a CSV file with 3D coordinates and identifiers. The cells were stained with Xenium to identify cell location as well as gene expression. Key cell types include cancer-associated fibroblasts, CD4^+^ T cells, and enterocytes. Users can zoom from the overall tissue structure down to individual cells and molecular signals, where the presence of RNA molecules indicates cell identity and activity. Regions with dense molecular signals may suggest proliferative, potentially pre-cancerous cells.These datasets are synthesized into an integrated model that represents an approximately thousand-fold difference in scale, comparable to the difference between a basketball and a grain of sand. This integrated 3D approach enables analysis of cell interactions and spatial patterns for offering insights into early cancer development and the organization of the tissue microenvironment. This scene features 18,285 cells on layer 16 with 20,919 surrounding RNA molecules in 3D. A detailed view is available in **Fig. 8** and **Fig. 2.f**.

**Fig. 8.**
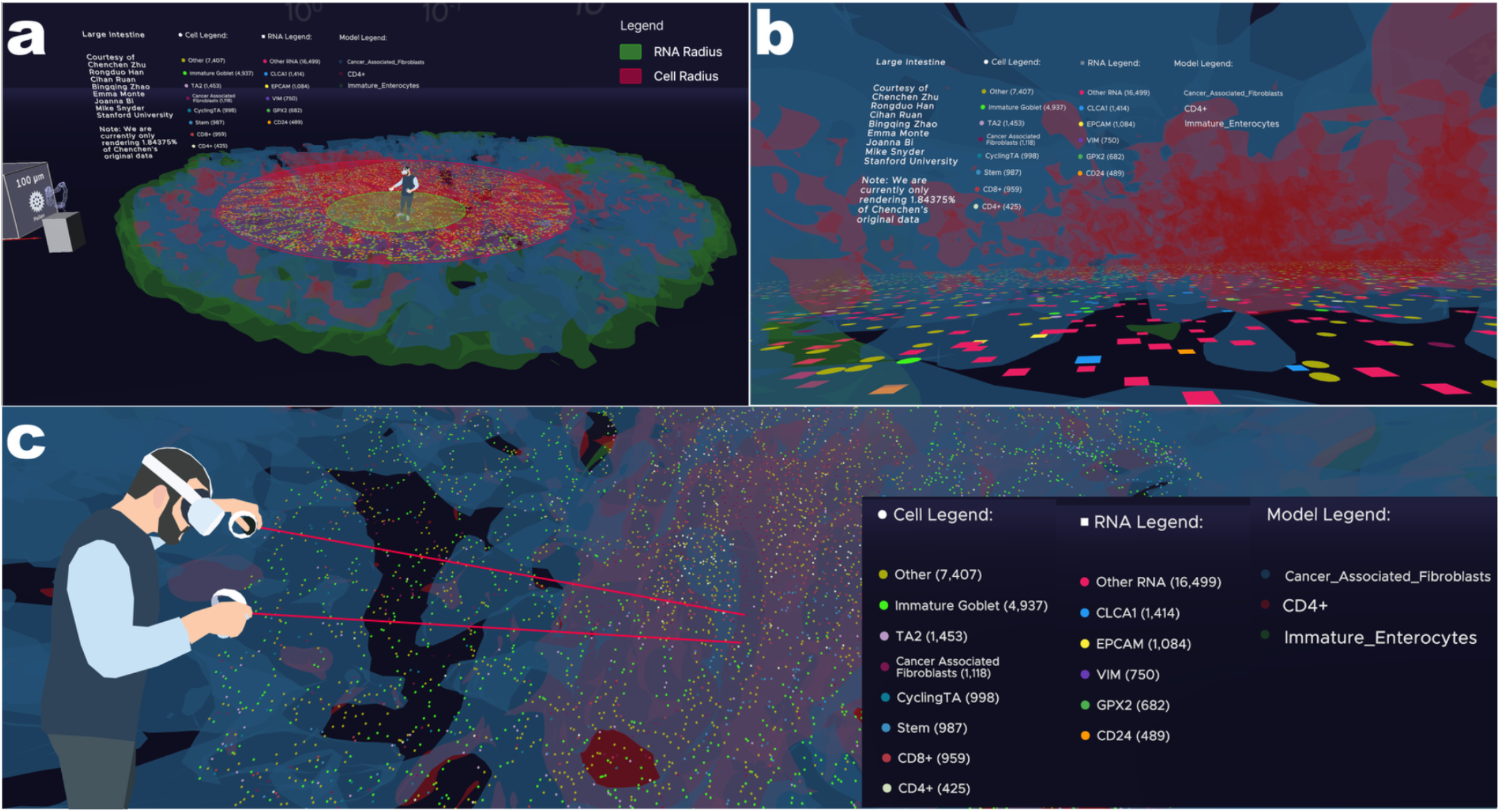
Snippets from the hubmap-large_intestine-zhu-100_microns-10_4 scene. **a**, A user in the combined large intestine data. Note the differing radii for cells and RNA in support of performance in VR as well as the similarity to the typical rendering of a disk-shaped galaxy, such as the Milky Way. **b**, A low-angle view of cells and RNA within the reconstructed 3D model. **c**, A user pointing towards cells and RNA molecules with the ray interactor on their VR controllers.

### cifar-liver-bader_xing-100_microns-10_4

*(allow 3D exploration and validation of liver data)*: This scene features human liver tissue and highlights the 3D architecture of the periportal region using SBF-SEM at nanometer resolution^34^. Shown are 3 anatomical landmarks: the portal vein, the bile duct, and hepatocyte organelle structures (on a subcellular scale) with mitochondria, cell nuclei, plasma membranes, and the endoplasmic reticulum. Approximately fifty hepatocytes are segmented, each enclosed by plasma membranes that define distinct cellular compartments. The user can grab, move, rotate, and scale each 3D model. The models were preprocessed using Blender (www.blender.org), an open-source 3D modeling tool, and Microscopy Nodes^35^ (extensions.blender.org/add-ons/microscopynodes). A detailed view is available in **Supplementary Fig. 5 and Fig. 2.d-e**.

### hra-multiscale-comparison-codex-100_microns-10_4

*(allow scale-accurate comparison of datasets from lymph node and small intestine in 3D space)*: Most scenes in the HRA Organ Gallery focus on 1 dataset (or a collection thereof) from 1 organ. When entering this scene, however, the user sees 4 entities with their true scale differences: the small intestine dataset HBM573.SWGH.988 from **hubmap-small_intestine-miao-100_microns-10_4**, the lymph node dataset SNT832.RKFV.643 from **sennet-lymph_node-enninful_farzad-1_centimeter-10_2**, the HRA 3D reference organ for the male right kidney, and the HRA 3D reference organ for the male skin from **hra-whole_body-hra-1_meter-10_0**. Additionally shown is a 3D model of the tallest building on Earth (Burj Khalifa, 828 meters).

These entities coexist in a shared space, see **Supplementary Fig. 6**. Typically, in each scene, datasets are scaled to about 1 meter in physical space. In this scene, however, the 2 datasets and the 3D reference organ are shown with the correct scale difference, where the small intestine data is 1 meter wide (see **Supplementary Fig. 6.d**), the lymph node data is 100 meters wide (two powers of 10 larger, see **Supplementary Fig. 6.c**), and the human body is 10,000 meters tall (four powers of 10 larger, see **Supplementary Fig. 6.a-b**). The 3D model of Burj Khalifa, however, is static and displayed in its real physical scale (828 meters); to illustrate this, it is rendered in red. As a result, when the scene starts, the skin 3D reference organ is already barely visible. For the small intestine and lymph node data, cell type groupings include immune cells, mesenchymal cells, and endothelial cells, which highlights shared biological components across tissues despite scale differences. Re-using existing work^36^, data preprocessing aggregated similar cell types into higher level categories, enabling clearer cross tissue comparisons. The user can dynamically resize the small intestine dataset and the lymph node and skin will scale concurrently, while Burj Khalifa provides a static, consistent large-scale benchmark (see see **Supplementary Fig. 6.b**). This multiscale visualization demonstrates how biological structures of vastly different sizes can be compared within a unified spatial framework. A detailed view is available in **Supplementary Fig. 6**.

### Adding new datasets

A SOP^12^ details how new datasets can be harmonized into the HRA Organ Gallery. It outlines 2 data categories: 3D models (e.g., OBJ, FBX, or glTF from v-CyCIF^9^ or and SBF-SEM^11^, OME-TIFF via STL, OME-ZARR) and data points (CSV, XSLX from CODEX^8^, Xenium^10^, Visium [www.10xgenomics.com/platforms/visium], or SWC^37^) and assigns responsibilities to predetermined roles in the process: the Data Provider (who contributes the data), the Designer (who ensures aesthetic fit), the Developer (who implements new scenes), and the Organ Gallery Lead (who coordinates all). The **Methods > Validation via SOP section** provides details regarding roles, procedures, and data formats.

## Discussion

HRA: Powers of Ten demonstrates that immersive, embodied 3D navigation can make multiscale biomedical data more interpretable via the HRA CCF than conventional 2D UIs allow. It lets Data Providers harmonize datasets from fundamentally different assays (CODEX, Visium, v-CyCIF, Xenium, SBF-SEM) into a shared spatial language and enables users to traverse the resulting scenes as a single continuous journey from the whole body down to subcellular structures. The Multiscale Elevator System’s central conceit (that each order of magnitude change in scale corresponds to a discrete floor) gives users an intuitive and consistent mental model for a kind of scale traversal that is otherwise difficult to convey. The biological insights reported across the 8 scenes (spatial organization of senescent cells in lymph node and brain tissue, layered cellular neighborhoods in the small intestine, 3D nerve-immune cell layout and cells with RNA molecules in the large intestine, as well as anatomical structures in the liver) suggest that the value of VR here is not merely illustrative. Features such as brushing and linking, exploded views, manual manipulation, rotation, and scaling of cells, embedded 2D visualizations in 3D environments, and cross-scale selection cones let users ask spatial questions (e.g., how are cell types distributed, what are the features of cells bordering a nerve, which senescence hallmarks occupy the same space, and how do cell types aggregate into neighborhoods, communities, and tissue units) that are harder to answer in 2D, so the platform can function as an analytical rather than only a presentation tool.

At the same time, we must acknowledge several limitations. The insights described are illustrative and qualitative; the paper does not report controlled comparisons of task performance, accuracy, or time-to-insight between the VR interface and standard 2D tools. The 8 scenes, while covering 5 organs and 4 scales, represent a small sample of available HRA data, and the datasets vary widely in size and preprocessing pipeline, which may limit generalizability of the workflow to other tissues, disease states, or assay types not yet represented. Hardware dependence on the Meta Quest platform, while broadening reach to over 20 million existing headset owners, also constrains adoption relative to institutions without VR infrastructure, and ergonomics concerns (motion sickness, headset fatigue) may arise. Scene creation currently depends on close collaboration between Data Providers, Designers, Developers, and the Organ Gallery Lead, as formalized in the accompanying SOP^12^; while this division of labor supported successful onboarding of ESIs in this study, the effort and specialized 3D modeling skill required (e.g., Blender-based preprocessing for SBF-SEM data) can still be a barrier to scaling data integration beyond expert teams.

Looking forward, our interest in applying this approach to more assay types and organs points toward VR’s most promising role: not replacing quantitative analysis but complementing it by surfacing spatial patterns worth testing computationally. Future work featuring immersive exploration and formal usability studies will expand the range of integrated organs and assays and lower the technical barrier to adding new datasets, which we hope will strengthen the case that multiscale VR studies like HRA: Powers of Ten can become standard infrastructure for building reference atlases rather than a specialized add-on.

## Methods

### Validation via SOP

The SOP^12^ entitled “Adding New Datasets to the Human Reference Atlas Organ Gallery” outlines the process for submitting 2D and 3D datasets for harmonization and visualization with the HRA Organ Gallery. It uses Jupyter Notebooks, Python, and Blender. The primary audience is Data Providers. The HRA Organ Gallery was developed with Unity 2022.3 (unity.com), a 3D real-time development platform. **Supplementary Notes 2-3** contain more information about native and custom Unity functionality used during development.

#### Roles and responsibilities

4 key roles govern the workflow. The **Data Provider** provides raw datasets and domain knowledge. The **Organ Gallery Lead** oversees the development of the HRA Organ Gallery and serves as the primary point of contact. The **Designer** creates VR scene specifications based on discussions between the Data Provider and the Organ Gallery Lead. Finally, the **Unity Developer** processes the submitted data and builds finalized scenes based on the Designer’s specifications under guidance from the Organ Gallery Lead. The Data Provider approves the final build of the scene.

#### Engagement

Typically, the Organ Gallery Lead initiates the collaboration by contacting the Data Provider via email. Upon engagement, the Organ Gallery Lead assigns the Data Provider a limited-access Google Drive folder for secure file uploads and a dedicated Acknowledgements document for capturing metadata, special handling requirements, and attribution information. Data Providers are expected to complete all required fields in this document before their scene is finalized and released.

#### Data formatting requirements

The SOP specifies formatting requirements depending on data type. For cell location and annotation data derived from CODEX^8^, Xenium^10^, and v-CyCIF^9^, datasets must be structured with columns for x, y, and optionally z coordinates, alongside a cell type or biological entity column. Preferred submission formats are 3D models (e.g., OBJ, FBX, or glTF from v-CyCIF or and SBF-SEM^11^, OME-TIFF via STL, OME-ZARR) and data points (CSV, XSLX from CODEX, Xenium, Visium [www.10xgenomics.com/platforms/visium], quantification tables for v-CyCIF, or SWC^37^). Blender is used for pre-processing if needed; otherwise, these files can be used in Unity right away. Custom processing for OME-ZARR is also completed and documented in the SOP^12^ but not featured in the version of the HRA Organ Gallery documented in this paper.

### Shared features between scenes

Each scene has a core set of features to enable multiscale navigation and exploration: an *Elevator Panel*, a *Scale Cube*, a *Skybox and Icon*, and an *Extraction Site Component*. **Fig. 3** shows all 4 shared features together.

### Elevator Panels

Every scene contains an Elevator Panel (see **Fig. 3.d)**, which allows the user to access all other scenes via buttons; each scene has a level number on which it is placed (expressed as a power of ten) and a name. If there are multiple scenes on the same level (such as on 10^−4^ = 100 microns), the level number is only displayed once. If a level has no scenes in it yet, it is colored white (locked), e.g., as of September 2026, the 10^−6^ level. If a scene can be entered, its button is colored HRA Blue (#201E3D). If it is active, i.e., if the user is currently in it, it is colored HRA Red (#FF0043). Both colors come from the HRA Style Guide^38^ and HRA Design System (docs.humanatlas.io/dev/design-system). At the very top of the Elevator Panel, a toggle button lets the user turn Elevator Transition Scenes on and off (see **Elevator Transition Scenes** section below). The Elevator Panel is instantiated in all scenes.

### Scale Cubes

While the scale of the virtual environment changes depending on the level the user travels to, the scale of the physical environment that the user is physically located in is, of course, persistent. As a result, the mapping between virtual and physical environments changes. To maintain a reference to the physical environment, a Scale Cube (see **Fig. 3.b**) with a volume of 1 cubic meter enables the user to get a visual reference for the length of 1 meter in the physical environment while in the virtual environment. On its user-facing side, the Scale Cube displays the current size that 1 meter in the physical environment corresponds to in this scene (e.g., 100 μm in **Fig. 3.b**) as well as the Icon for that level (e.g., a pollen, see **Skyboxes and Icons** section below) and the name of the object that the Icons depicts (e.g., “Pollen”). The object on the Icon represents everyday objects that are understandable without domain expertise. The Scale Cube auto-rotates around its y-axis to always face the user.

### Skyboxes and Icons

Skyboxes (see **Fig. 3.a** and **Supplementary Fig. 1)** are large-scale textures (typically cubemaps) commonly used in 3D environments that simulate a sky environment by wrapping around an entire scene (docs.unity3d.com/2022.3/Documentation/Manual/skyboxes-using.html). In the HRA Organ Gallery, Skyboxes communicate scale transitions and to support user orientation during navigation across biological levels. Their primary function is to serve as large-scale environmental cues to reduce visual emptiness and help users contextualize their relative size in each level. The conceptual foundation for the Skyboxes is the Nikon Universcale (www.nikon.com/company/corporate/sp/universcale) and the “Powers of Ten” movie ^20^.

All Skybox illustrations were designed in Figma (www.figma.com). The design approach emphasized clean vector shapes to maintain legibility at large viewing distances. Colors were derived from the HRA Style Guide ^38^ and HRA Design System (docs.humanatlas.io/dev/design-system) for stylistic continuity across light and dark modes. Subtle luminance gradients were introduced to create a sense of atmospheric depth and to avoid the flat appearance typical of uniform backgrounds. Each Skybox was produced at high resolution to reduce stretching and pixelation when mapped to a spherical projection. The vertical placement of objects above the horizon was also determined through trial and error. Object height was tested both in the spherical Skybox in VR and in the flat 2D rectangular layout in Figma to ensure the object appeared at the intended position once projected. The technical development of the Skyboxes required extensive iteration. Initial tests revealed significant distortion when flat 2D compositions were wrapped onto a spherical Skybox surface in Unity. To resolve this, projection centers and object placements were recalibrated, and illustrations were repositioned using adjusted polar coordinates. Early tests also revealed color discrepancies between PNG exports and their appearance in VR due to scene lighting. Converting Figma assets from PNG to OpenEXR (openexr.com/en/latest) significantly improved visual accuracy under high dynamic range (HDR, www.adobe.com/creativecloud/photography/discover/hdr.html), which ensured more reliable tone mapping. A light and a dark version of all the Icons were also created to test what works best for our purpose. The final set consists of eleven Skyboxes exported as OpenEXR textures for optimal HDR compatibility. These were integrated into Unity as non-interactive environmental layers with optimized rendering settings to improve performance across standalone Meta Quest headsets.

### Extraction Site Component

To indicate the spatial origin of a dataset in the human body, each scene features an Extraction Site Component (see **Fig. 3.c** with the extraction site for **hubmap-small_intestine-miao-100_microns-10_4** in the duodenum of the small intestine), which shows where a dataset is located with an HRA 3D reference organ, enabled by prior registration of the spatial origin via the RUI. This component imitates the look of the RUI and Exploration User Interface (EUI)^4^ but introduces a district visual style to fit the visual style of the HRA Organ Gallery. While organ models in the HRA typically have materials that aim to imitate their physical look, 3D reference organs in Extraction Site Components are assigned a material that uses a Fresnel shader with transparency and a dark blue albedo color. An overlaid extraction site marker (cuboid) with a red material is always rendered on top, even if located deep within an organ, to indicate the position, rotation, and scale of the extraction site of the dataset. New Extraction Site Components can be created via a Unity Editor script (learn.unity.com/tutorial/editor-scripting), which needs pointers to a 3D reference organ, a cached HRA API response as a *ScriptableObject* (docs.unity3d.com/2022.3/Documentation/ScriptReference/ScriptableObject.html), and a tissue block/sample ID. Details on these are provided in **Supplementary Note 2**.

## Acknowledgments

In each scene, 2D text panels acknowledge the Data Providers and all relevant metadata plus a verbose description of the data presented.

## Shared scripts and Unity features between scenes

Scenes were built in [Unity](https://unity.com) 2022.3 with XR Interaction Toolkit 2.6. Data such as cell positions, color maps and cached HRA API responses are stored as [ScriptableObjects](https://docs.unity3d.com/2022.3/Documentation/Manual/class-ScriptableObject.html) so the application runs offline (see **Supplementary Note 2**). Scenes share a set of custom components for visualizing cells, building legends and highlighting cell types across the scene (brushing and linking). Two Editor tools prepare data before the application is built: *IngestCellPositions* converts CSV files of cell positions and types into *ScriptableObjects*, and *IngestScene* caches 3D reference organs and tissue blocks from the HRA API before each HRA release (see **Supplementary Note 3**).

### Elevator Transition Scenes

The HRA Organ Gallery’s Multiscale Elevator System (see **Supplementary Fig. 2**) includes a cylindrical elevator that allows users to travel between different biological scales, ranging from 10^0^ (whole body level) to 10^−9^ (molecular level). Each Elevator Transition Scene lasts 30 seconds and contains either a human body model for reference (for scales 10^0^ through 10^−6^) or a group of red blood cells in a blood vessel (for 10^−7^ and below). During the scene, the user is placed inside the elevator and moves either down or up, depending on whether they are traveling to a smaller or larger scale. After 30 seconds, the scene routes the user to the selected destination scene. Elevator Transition Scenes can be skipped via a toggle on the Elevator Panel (see red button in the top row of the Elevator Panel in **Fig. 3.d**). A demonstration video is available at cns-iu.github.io/hra-organ-gallery-supporting-information/#videos.

Two major challenges had to be solved to make Elevator Transition Scenes work: relative size and relative speed.

#### Relative size

In each Elevator Transition Scene, the sizes of both the user in the elevator and the reference object (human or red blood cells) are adjusted such that their scale difference corresponds to the level to which the user travels. For example, if the user travels to a scene on the 10^−4^ level, the human body will be 4 orders of magnitude larger than it would typically appear. Starting at the 10^−7^ level, however, the scale difference (which roughly corresponds to the distance between Chicago, IL, and Cairo, Egypt) becomes so large that the human body becomes unrecognizable and invisible to the user due to the large draw distance. As a result, for the 10^−7^ level and below, a different reference object is used (red blood cells). **Supplementary Fig. 2** shows these 2 reference objects from different angles (**a-b** for the human, **c-d** for the red blood cells). Moreover, **Supplementary Fig. 2.b and d** feature the user’s view from inside the elevator while looking at the adult human and a group of red blood cells, respectively.

#### Relative speed

A related issue arises when the scale difference between user and reference object becomes increasingly large. To give the user the impression of moving, the relative position between the elevator and the reference object needs to change perceptably. However, at vast scale differences, even fast movement (greater than 100 meters per second) is not sufficient to induce this feeling of motion. Additionally, fast movement can introduce rendering challenges for Unity. To address this issue, a particle system was implemented, where particles emerge from the bottom of the elevator in a circular pattern and move upward with visible trails, which creates the illusion of the elevator moving through the scene. This is also explained in the demonstration video at cns-iu.github.io/hra-organ-gallery-supporting-information/#videos.

### Preprocessing and scene setup

#### hra-whole_body-hra-1_meter-10_0

In this default start scene, the user is presented with a work area that includes a desk (see **Supplementary Fig. 3.a**). In the scene, they can move, rotate, and scale all 81 3D reference organs of the HRA v2.5 (humanatlas.io/3d-reference-library?version=11&organ=All%20organs) as well as tissue blocks registered into them. When loaded, the user is presented with an organ selection keyboard (see **Supplementary Fig. 3.b**), where they can pick an organ to inspect. Organ selection buttons consist of a base button model with a symbol on top of it using HRA organ icons, see HRA Style Guide^38^ and HRA Design System (docs.humanatlas.io/dev/design-system), and the name of the organ below it. The top row features functionality buttons to reset the organ position, rotation, and scale, a sex selection button (if applicable), a laterality selection button (if applicable), and a button to hide tissue blocks. The latter allows the user to inspect organs whose details are obstructed by overlapping tissue blocks. Buttons to switch between user movement and the user’s ability to “explode” tissue blocks as well as to show cell type populations are present but inactive and will be made active in future releases.

Each button has 3 possible states: ready (button can be pressed), pressed (button is pressed), and locked (button cannot be pressed). Colors from the HRA Design System are assigned based on state. Button states are handled by a C# script that captures user input with events-based C# architecture (learn.microsoft.com/en-us/dotnet/csharp/programming-guide/events) and checks which organ, sex, and laterality combinations are possible. When an organ button is clicked, the organ is loaded into a grab cylinder (e.g., the female brain in **Supplementary Fig. 3.c**) so it can be moved, rotated, and scaled for anatomical exploration (see the scaled up placenta and umbilical cord in **Fig. 2.a**). The loaded organ has all its tissue blocks, which are retrieved via the HRA API endpoints at apps.humanatlas.io/api#get-/v1/reference-organ-scene and apps.humanatlas.io/api#get-/v1/scene. These API calls are made during development using Editor scripts, and the response is cached as a *ScriptableObject* in Unity, i.e., a data container with a matching C# class that extends the *ScriptableObject* class in the Unity code base (see **Supplementary Note 2**). This allows the application to be updated using the most recent HRA data but without the need for an internet connection at runtime. The code at github.com/cns-iu/hra-organ-gallery-in-vr-preprocessing/tree/main/preprocessing-glb-download is used to download all available 3D reference organs of the most recent HRA release at the time of running the code. Once an organ is loaded in the grab cylinder, it is highlighted inside the whole body of either the male or female body on the table (see **Supplementary Fig. 3.d**). In addition to the Scale Cube in the back of the table, a smaller scale cube (one order of magnitude smaller) is shown next to the user (see **Supplementary Fig. 3.e**). It scales up and down if the user scales the selected organ up and down.

The position, rotation, and scale transformation for this scene are prepared with the *IngestScene* Editor script calling the HRA API (apps.humanatlas.io/api), see **Supplementary Note 3**.

#### sennet-lymph_node-enninful_farzad-1_centimeter-10_2

Like other scenes in HRA: Powers of Ten, the CODEX data for the lymph node in this scene is visualized with the *Visualizer3D* component, which is derived from a *VisualizerBase* component that defines base functionality to take a list of cell types in 2D or 3D, lays it out in space, and applies formatting as needed with brush-and-link functionality (see **Supplementary Fig. 4** and implementation details in **Supplementary Note 3**).

#### sennet-brain-phatnani-1_millimeter-10_3

This scene enables visual exploration of 8 senescence hallmarks across 4,992 spots in Visium data from the middle frontal gyrus of the brain of a 74-year-old female, human donor. Relevant SenNet IDs are: SNT239.DNSH.749 (donor), SNT326.PVLT.534 (tissue block), SNT873.QHZM.365 (dataset), and SNT956.RQLW.577 (upload). The scene features 2 major items: a 3D spike plot to visualize expression values for 8 color-coded senescence hallmarks (see **Fig. 5.a**) and a gallery of 10 2D plots made with *plotly* (plot.ly, see **Fig. 5.b**) that were imported as PNG textures, then applied to 2D surfaces in the scene. The user can grab, rotate, and bring these 2D plots with them when stepping into the 3D spike plot.

##### 3D spike plot

Each dot contains average expressions for 8 senescence hallmarks for all the cells captured in that circular area. If a 3D spike goes above the plane of the cell spot, the corresponding senescence hallmark is positively expressed and below the plane of the spot if it is negatively expressed (see **Fig. 5.a and c**). The 3D spike plot was created using standard UI components in Unity, i.e., dot sprites for 4,992 cell spots on the ground (docs.unity3d.com/2022.3/Documentation/ScriptReference/Sprite.html) and line meshes for the 3D spikes (docs.unity3d.com/2022.3/Documentation/ScriptReference/Graphics.DrawMeshNow.html). To avoid visual clutter in the 3D spike plot, lines are only shown for cell spots that have a low mean and a high variance among their senescence hallmark expressions. As a result, only 1,576 lines are drawn for 197 cell spots; if all were shown, the user would have to navigate through 39,936 lines for 4,992 cell spots.

##### 2D plots

While the 3D spike plot enables visual and spatial exploration of 8 senescence hallmarks in 3D space concurrently, the gallery of 2D plots presents scatter graphs where color intensity encodes the expression value for exactly 1 hallmark in each plot. 2 additional scatter graphs show which cell spots get senescence biomarkers and where those cell spots are in the Visium grid (see **Fig. 5.d**).

#### hubmap-large_intestine-wong-100_microns-10_4

Two major datasets in v-CyCIF^9^ are explorable in this scene: a light-sheet microscopy dataset and a healthy confocal dataset, both from the large intestine (see **Fig. 6.a**). They are color-coded and share the same color scheme. A legend is shown to the user in the scene. Major pre-processing was done to make the CD8 cells interactable, make visual encoding adjustable by the user, and enable scaling of the light-sheet dataset. The adjacent healthy large intestine model was provided by co-author Wong in 4 separate OBJ files representing different biological components: CD8, CD31, Claudin-7 and TUBB3. These 4 files together form a single combined structure with significant spatial overlap and interconnection between layers.

For the light-sheet crop (**Fig. 6.a**, left), co-author Wong provided a quantification table (CSV) with cell IDs, positions and volumes, and a 3D model of CD8 cells (OBJ) whose objects had generic names (e.g., “CD8.001”). Because manual mapping was impractical for this many cells, a Python script in Blender matched each 3D cell to its table row. It first aligned the two coordinate systems using a reference cell identifiable in both, then accepted a match only if position (≤3.0 scene units) and volume (≤20% difference) both agreed. The Data Provider validated the resulting mapping (see **Supplementary Note 4**).

Cells can be grabbed, inspected, and returned to their original position, and colored by volume, sphericity, minimum intensity or distance to origin (see **Supplementary Note 5**).

##### Mesh optimization and performance considerations

When initially loaded, the combined light-sheet crop and the healthy confocal large intestine model contained approximately 6 million polygons, which was well above the recommended target of ~1.3-1.8 million polygons for VR scenes (see **Performance metrics for Meta Quest VR headsets** section). To reduce geometric complexity, all 4 OBJ files for the confocal dataset were imported into Blender and decimated individually with the *decimate* modifier (docs.blender.org/manual/en/latest/modeling/modifiers/generate/decimate.html). A decimation factor of 0.05 was selected through trial and error, which resulted in each mesh retaining approximately 5% of its original polygon count while maintaining acceptable visual fidelity. After decimation, the healthy large intestine model was imported into the existing Unity scene. With both the previously integrated CD8 model and the new large intestine model present, the total polygon count in the scene was reduced to approximately 3 million. Runtime testing showed that the scene continued to run smoothly, no significant frame rate drops were observed, and performance remained acceptable despite exceeding the recommended target polygon budget. During normal usage, not all of the polygons are viewable concurrently.

#### hubmap-small_intestine-miao-100_microns-10_4

Because this scene contains various features in 3D space, a “Stand Here” sign on the ground marks a user’s recommended initial position (see **Fig. 7.a**). 4 layers show the same cells in 4 different colors based on 4 different layers of aggregation: cell type, neighborhood, community, and tissue unit. With their right hand, the user can then place an adjustable, cross-layer selection cylinder to highlight the same cell on all 4 layers, with context provided by a text canvas that shows the number of selected cells per layer (see **Fig. 7.b** and **Fig. 2.c**). A trash can icon allows the user to delete all selection cylinders. Additionally, an alluvial diagram on the ground shows how many cells on which cell type, neighborhood, community, and tissue unit were selected by using color intensity (white to red) and numeric labels (see **Fig. 7.c**). Additionally, an Extraction Site Component highlights the spatial origin of the data in the proximal jejunum of the small intestine (see **Fig. 7.d**). The data^31^ comes from HuBMAP dataset HBM573.SWGH.988 in sample HBM787.FHZM.948 from donor HBM578.DXWB.873.

To implement the text canvas with the number of selected cell instances per layer, a *Canvas* (see **Supplementary Note 2**) was set up along with several scripts: *CellLogger*, a data processing script, and *CellTypeDisplay*, a script to print the uniquely identified cells per layer of aggregation across as a simple label. *CellLogger* processes the selected data labels into a dictionary grouped by x-y coordinates for each entity, while *CellTypeDisplay* prints the unique entities in the selection cylinder to a label in front of the visualization in the scene.

To create the alluvial diagram, several implementation steps were performed: A *GraphVisualizer* script uses the data grouped by x-y coordinates obtained from the *CellLogger* script to map relationships between the unique entity types into a graph. It also logs the number of instances of each entity in each entity node, and a weight along the graph edges showing the number of entity relationships of that type. The nodes and edges are also color coded to show a gradient of weights as they increase/decrease. The trash can icon was remapped to clear the alluvial diagram on click.

#### hubmap-large_intestine-zhu-100_microns-10_4

This level depicts an early-stage, cancer-afflicted large intestine section reconstructed as a 3D tissue map and overlayed with spatial cell type and RNA transcript data with the goal of enabling users to examine how disease-associated cellular and molecular features are spatially organized within the tissue. Traditional histological analysis relies largely on thin tissue sections, typically approximately 5 microns thick, in which few, if any, cells are captured in their entirety. As a result, 2D imaging often fails to reveal critical aspects of tissue architecture and cellular ultrastructure that become apparent only through 3D analysis. Using Space-map ^33^, these limitations were addressed by constructing a 3D tissue model and developing visualization approaches that enable exploration across multiple biological scales, from whole tissues and individual cells to molecular features such as RNA localization.

The **hubmap-large_intestine-zhu-100_microns-10_4** scene focuses on stem cells and their differentiated progeny within the colonic crypt-villus axis, which are essential for maintaining normal intestinal mucosal function, as well as on diverse immune cell populations residing in secondary lymphoid structures such as Peyer’s patches. 3D visualization and analysis of these tissue-resident immune cells enable detailed characterization of their spatial organization and interactions within their native microenvironments.

Co-author Zhu provided the 3D tissue reconstruction model and its cell layers plus RNA molecules. The primary objective of this scene was to integrate the spatial data visualization layers derived from co-author Zhu’s dataset directly within the corresponding 3D model. To facilitate exploration of the embedded data, the opacity of the 3D model was reduced to an alpha value of 0.2 to allow users to navigate through both the model and the visualization layers simultaneously. This design emphasizes immersive interaction by displaying only data points within a defined proximity to the user, regardless of their original layer assignment, as the user moves through the environment.

An Extraction Site Component in the scene marks the sigmoid colon as the anatomical structure where the tissue was extracted. The data comes from HuBMAP sample HBM896.ZPRW.239, which was retrieved from HuBMAP donor HBM694.QDFG.746.

An overview of this scene is presented in **Fig. 8**. This scene posed 2 major technical challenges:

##### Data integration

It contains a 3D model (OBJ), 29 layers of cells in 2D (CSV), and, for each of these layers, RNA molecules (CSV). These 3 data types had to be aligned in the same virtual space. The guiding principle for the visual style of this scene was the typical rendering of a disk-shaped galaxy, such as the Milky Way (see **Fig. 8.a**).

##### Performance

The original 3D model already had 1.6 million polygons. Each of the 29 cell layers contained around 80,000+ cells, which, if all 29 layers were rendered, would amount to 2.32 million cells in a singular scene, which would be too computationally expensive on the Meta Quest hardware. The scene exemplarily shows cell layer 16 (18,285 cells) with RNA layer 16 (20,919 RNA molecules) as these are roughly in the center of the z-axis (depth) of the 3D model. To address this challenge, the *decimate* modifier in Blender was used to reduce the polygon count by ~41% to 942,436. Additionally, a visibility radius around the user ensures that only cells and RNA molecules within 300 (cells) and 100 microns (RNA) are rendered, respectively, see **Fig. 8.a**. A detailed view of cells and surrounding RNA molecules is available in **Fig. 8.b**. An illustration of a user traversing the data alongside the legend is shown in **Fig. 8.c. Fig. 2.f** shows an alternative angle on a user exploring a local cluster of cells and RNA molecules.

##### Data preprocessing

Generating visualization layers required extensive preprocessing. While originally all 29 individual CSV files (with native 2D coordinates) were downloaded and given a z-position with an iterative z-axis offset of 0.03 meters in VR to fill out the depth of the 3D model, the deployed **hubmap-large_intestine-zhu-100_microns-10_4** focuses on cell layer 16 and RNA layer 16 based on co-author Zhu’s input. On these layers, cell and RNA molecule labels were aggregated to the top-5 plus an “other” category to reduce the number of colors used in the visualization (cells: Immature Goblet, TA2, Cancer Associated Fibroblasts, CyclingTA, Stem; RNA: CLCA1, EPCAM, VIM, GPX2, CD24). These 2 CSV files were then processed using the *IngestCellPositions* Editor script in Unity. This made the combined CSV file more easily usable at runtime and editable in the Unity Inspector. For this combined CSV file, the ingest process generated 2 *ScriptableObjects*: The first *ScriptableObject* stores the spatial coordinates (x, y, z-positions) of all cells across all layers with their corresponding cell type label. The second *ScriptableObject* contains summary statistics describing the frequency of each unique cell type per layer. These data structures serve as the primary inputs for subsequent visualization.

##### Visualization

3D visualizations are generated using the *Visualizer3D* component, which uses both these *ScriptableObjects* produced during data preprocessing (see **Supplementary Notes 2-3**). While rendering, the application instantiates individual *GameObjects* representing cells with *CellData* components and places them at spatial locations defined by the coordinate data stored within each layer’s *ScriptableObject*. Cell labels and frequencies are obtained from the corresponding *_frequency ScriptableObject*. The resulting visualizations are then positioned within the Unity scene using appropriate z-axis spacing to ensure accurate alignment with the reconstructed tissue model. As a result, the final scene combines the 3D tissue reconstruction model with its corresponding spatial cellular data. To accommodate the sheer volume of cells, the *IngestCellPositions Editor* script was set to ingest only 1/64^th^ of the original number of cells, making it comfortable to render 18,285 cells and 20,919 RNA molecules of layer 16 with the targeted hardware.

#### cifar-liver-bader_xing-100_microns-10_4

The data for this scene comes from the caudate lobe of the liver’s periportal region (see extraction site in **Supplementary Fig. 5.a**.). 3 models were shared and pre-processed (**Supplementary Fig. 5.b**). A detailed view is available for the portal vein (**Supplementary Fig. 5.c**), bile duct (**Fig. 2.d**), and hepatocyte organelle structures (**Fig. 2.e**).

##### Pre-processing

Co-authors Bader and Xing shared 3 TIFF files containing slice data of cross sections for portal vein, bile duct, and hepatocyte organelle structures. As received, they were not ready for VR, since they lacked mesh-based surfaces to render. The SOP for adding new datasets to the HRA Organ Gallery^12^ outlines explain 2 methods to generate 3D models (FBX files) from these TIFF files:

1. Use the Microscopy Nodes^35^ plug-in (extensions.blender.org/add-ons/microscopynodes) in Blender to load TIFF files, then export them as FBX files.
2. Process TIFF files into STL files using Fiji (imagej.net/software/fiji/), then convert STL to FBX in Blender and reduce polygon count with MeshLab (www.meshlab.net).

While method #1 is the preferred method to convert these TIFFs into FBX for usage in Unity, for this scene, it preserved too much of the surface definition in the FBX, which led to a surplus of polygons for the Meta Quest 3 VR headset (see **Performance metrics for Meta Quest VR headsets** section). While Blender’s *decimate* modifier can reduce polygons in 3D models, it is not effective for large models with many internal structures. To reduce polygon counts effectively while preserving most of the surface definition, method #2 was used.

##### Scene setup

An Extraction Site Component (see **Supplementary Fig. 5.a**) shows the spatial origin of the liver sample from which the TIFF files (and FBX files after conversion) were generated. A cuboid table displays the portal vein, bile duct, and hepatocyte organelle structures (see **Supplementary Fig. 5.b**). To enable interaction, an *XRGrabInteractable* was added to each of them for movement and rotation, along with a *XRGeneralGrabTransformer* to enable scaling (see **Supplementary Note 2**). For aesthetic reasons, the model materials were adjusted to appropriately different but muted colours that are aligned with the HRA Design System.

#### hra-multiscale-comparison-codex-100_microns-10_4

This scene is set up to convey to users how the same cell type distribution can exist in multiple biological entities (entire lymph node and proximal jejunum) at different scales using the CODEX data from **sennet-lymph_node-enninful_farzad-1_centimeter-10_2** and **hubmap-small_intestine-miao-100_microns-10_4** (see **Supplementary Fig. 6**). Since the main focus of this scene was to visualize common cell types across 2 scales, a common cell type color map was needed.

Related HRA work^36^ on crosswalking and aggregating cell to higher-level cell types using Cell Ontology (CL)^39^ was used; the CSV file with these higher-level cell types is available on GitHub (github.com/cns-iu/hra-cell-distance-analysis/blob/1f73eed7032af63a6174b52bd0a8057e38c051df/data/mapping_files/generated_cell_type_complete_crosswalk.csv).

To facilitate comparison across shared, higher-level cell types, the following steps were needed:

1. The original CSV files with 3D cell positions and types were enriched with additional columns to represent level 3 (original, lowest aggregation), level 2 (medium aggregation), and level 1 (highest aggregation) cell types.
2. A mapping from cell type to color was created to standardize colors across the common cell types. This was done via an Editor script. A *ScriptableObject* was used to make the color map available at runtime and editable in the Unity Editor (see **Supplementary Note 2** for technical details).
3. Both datasets were visualized in the scene, but the width of the lymph node was increased by 2 orders of magnitude from the user’s perspective, because it is 2 orders of magnitude larger than the slice of proximal jejunum. In their processed form, both datasets share the same color mapping.
4. The HRA 3D reference organ for the male skin (humanatlas.io/3d-reference-library?version=11&organ=Skin) was placed in the scene with the correct scaling difference to the lymph node.
5. Finally, to convey the scale difference between the 2 biological data visualizations and the 3D reference organ with regards to a known real-world object, a 3D model of Burj Khalifa, the tallest building on earth as of September 2026 at a height of 828 meters, was added to the scene in its true (non-scaled) size; a flat plane attached to its summit served as visual reference for its height of 828 meters. This model does not change scale.
6. Next, *XRGrabInteractables* and *XRGeneralGrabTransformers* (see **Supplementary Note 3**) were added to both visualizations to allow the user to scale the spatially arranged data visualization. A scale synchronizer script on an empty *GameObject* synchronizes the scale of the proximal jejunum data, the lymph node and the HRA 3D reference organ for the male skin, to show how they scale uniformly together.
7. Finally, a reset button next to the cell type legends enables users to reset the visualizations to their position, rotation, and scale.

#### Performance metrics for Meta Quest VR headsets

As of September 2026, Meta provides performance targets for testing and analysis during development for Meta Quest VR headsets at developers.meta.com/horizon/documentation/unity/unity-perf. This comes in the form of recommended draw calls (i.e., commands sent from the CPU to the graphics API requesting that the GPU draw an object on the screen) given light, medium, or busy simulations (i.e., the number of active graphical elements in the scene). While no concrete metrics are provided for what constitutes light, medium, or busy simulations, it can reasonably be assumed that, despite the high-quality 3D data, the scenes presented in this paper constitute at most medium simulations due to the low number of 3D objects on screen, the simple environment, the lack of background elements and animations, the use of baked or mixed lighting instead of real-time lighting, and the lack of post-processing effects. Meta recommends 400-600 draw calls maximum for these scenes. Likewise, in terms of polygon counts, for the Meta Quest 3 and 3S, 1.3-1.8 million in the user’s field of view are recommended.

## Data availability

All data needed to evaluate the conclusions in the paper are present in the paper and/or the Supplementary Information and on GitHub. A companion website is available at cns-iu.github.io/hra-organ-gallery-supporting-information. To improve documentation and enable readers without VR headsets to learn about the scenes in HRA: Powers of Ten (incl. one documenting Elevator Transition Scenes), videos were compiled at cns-iu.github.io/hra-organ-gallery-supporting-information/#videos, which feature narration by the Data Provider(s) and the Organ Gallery Lead in a recorded VR demo. A workshop to jumpstart pilot project integration was held at the Bioinformatics and Computational Biosciences Branch at National Institute of Allergy and Infectious Diseases and is documented at cns-iu.github.io/workshops/2024-10-24-jumpstart-workshop. Supporting information for this paper is available at github.com/cns-iu/hra-organ-gallery-supporting-information.

81 3D reference organs of HRA v2.5 are available at humanatlas.io/3d-reference-library and 3d.nih.gov/collections/hra. Datasets SNT873.QHZM.365 and SNT832.RKFV.643 with their entire provenance are available at data.sennetconsortium.org. Datasets HBM573.SWGH.988 and sample HBM896.ZPRW.239 with their entire provenance are available at portal.hubmapconsortium.org.

The model of Burj Khalifa for **hra-multiscale-comparison-codex-100_microns-10_4** was downloaded from sketchfab.com/3d-models/free-burj-khalifa-dubai-c1d6f5884c9c4a56b8d8f9c5555f1902; it was made with Blender 2.90.

The application is available for Meta Quest 2, 3, 3S, and Pro headsets on the Meta Store: www.meta.com/en-gb/experiences/hra-organ-gallery/5696814507101529. All code is accessible at github.com/cns-iu/hra-organ-gallery-in-vr. Preprocessing code for this paper is available at github.com/cns-iu/hra-organ-gallery-in-vr-preprocessing.

## Author contributions

CRediT: Conceptualization: A.B., K.B.; Data curation: A.B., C.Z., A.Y.H.W.,, A.E., Y.M., N.F., B.W.H., M.P., C.M., N.S., J.M., C.X., Y.J.; Formal Analysis: C.Z., A.Y.H.W.,, A.E., Y.M., N.F.; Funding acquisition: A.B., J.W.H., R.F., P.K.S., M.S., K.B., G.D.B., H.P., V.M.; Investigation: A.B.; Methodology: A.B., Y.K., K.P., S.C.; Project administration: A.B.; Resources: B.W.H., K.B., Y.J.; Software: A.B., B.W.H., Y.K., K.P., S.C.; Supervision: A.B., J.W.H., R.F., P.K.S., M.S., K.B., G.D.B., H.P., V.M.; Validation: A.B., C.Z., A.Y.H.W.,, A.E., Y.M., N.F., M.P., C.M., N.S., J.M., C.X.; Visualization: A.B., J.K., Y.K., K.P., U.P., S.C.; Writing – original draft: A.B., J.K., Y.K., K.P., S.C.; Writing – review & editing: A.B., C.Z., A.Y.H.W.,, A.E., Y.M., N.F., J.K., Y.K., K.P., K.B., S.C., M.P., C.M., N.S., J.M., C.X., P.L.

## Acknowledgments

We would like to extend our deep gratitude to Darrell Hurt, Ph.D., Meghan McCarthy, Ph.D., Phil Cruz, Ph.D., Kristen Browne, M.Sc., and Victor Starr Kramer from the Bioinformatics and Computational Biosciences Branch at the National Institute of Allergy and Infectious Diseases for co-hosting the HRA: Powers of Ten Workshop on October 24-25, 2024 (cns-iu.github.io/workshops/2024-10-24-jumpstart-workshop) and for providing resources in the form of hardware and developer time. We would further like to thank Niteesha Jangam for helping us organize the workshop and Danial Qaurooni for creating and managing extraction sites for HuBMAP, SenNet, and other efforts.

The HRA (humanatlas.io) is under active development by HuBMAP, SenNet, the Kidney Precision Medicine Project, GenitoUrinary Developmental Molecular Anatomy Project, and the National Institute of Diabetes and Digestive and Kidney Diseases with expert input by the HRA Editorial Board (humanatlas.io/about#editorial-board) and in close collaboration with experts from 20+ other consortia.

This research has been supported by the following awards:

- The NIH Common Fund through the Office of Strategic Coordination/Office of the NIH Director:

- HuBMAP:
  ▪ 3OT2OD026671 (A.B., B.W.H., U.P., K.B.)
  ▪ 3OT2OD033756 (A.B., B.W.H., J.K., Y.K., K.P., U.P., Y.J., K.B.)
  ▪ U54HG010426 (C.Z., M.P.S.)
  ▪ U54HG012723 (C.Z., M.P.S.)
  ▪ OT2OD026675 (A.B. as NIH JumpStart Award 2023)
  ▪ 3OT2OD033759 (A.B. as NIH JumpStart Fellowship 2024, J.K., Y.K., K.P., S.C.)
  ▪ 3OT2OD033759-01S3 (A.Y.H.W., C.Z., as NIH JumpStart Fellowship 2024)
  ▪ 3OT2OD033759-01S4 (Y.M., J.W.H.)
- SenNet:
  ▪ U24CA268108 (A.B., B.W.H., U.P., Y.J., K.B.)
  ▪ U54AG076043 (A.B., A.E., N.F., Y.J., R.F.)
  ▪ U54AG079759 (A.E., N.F., R.F.)
  ▪ U54AG076040 (M.P., C.M., N.S., H.P., V.M.)
  ▪ 3U54AG075936 (Y.M., J.W.H.)
- Common Fund Data Ecosystem (CFDE):
  ▪ OT2OD030545 (A.B., B.W.H., K.B.)
  ▪ 1R03OD039970-01 (A.B., B.W.H., J.K., Y.K., K.P., S.C., U.P.)
- National Institute of Diabetes and Digestive and Kidney Diseases:
  - Kidney Precision Medicine Project: U01DK133090 (A.B., B.W.H., K.B.)
  - U2CDK114886 (A.B., B.W.H., K.B.)
  - U24DK135157 (B.W.H., K.B.);
- National Cancer Institute (NCI):
  - UG3CA257393 (A.E., N.F., R.F.)
  - UH3CA257393 (A.E., N.F., R.F.)
  - R01CA245313 (R.F.)
  - U54CA274509 (R.F.)
  - U54CA268083 (R.F.)
  - U01CA294514 (R.F.)
  - U54-CA268072 (A.Y.H.W., P.K.S.)
- National Institute of Mental Health (NIMH):
  - RF1MH128876 (R.F.)
  - RM1MH132648 (R.F.)
- This research is in part based on work supported by a CIFAR catalyst award (A.B., Y.K., J.K., C.X., Y.J., G.D.B., K.B.).
- K.B. and G.D.B. are co-directors of and are funded by the MacMillan Multiscale Human program by CIFAR.
- K.B. is also supported via a Stiftung Charité Visiting Fellowship via Berlin Institute of Health at Charité (BIH).

The funders had no role in study design, data collection and analysis, decision to publish, or preparation of the manuscript. The content is solely the responsibility of the authors and does not necessarily represent the official views of the NIH.

This research uses IU’s computing infrastructure which was supported in part by Lilly Endowment, Inc., through its support for the Indiana University Pervasive Technology Institute.

## Additional information

### Competing interests statement

G.D.B. advises BioRender.

R.F. is scientific founder and adviser for IsoPlexis, Singleron Biotechnologies, and AtlasXomics. The interests of R.F. were reviewed and managed by Yale University Provost’s Office in accordance with the University’s conflict of interest policies.

P.K.S. is a cofounder and member of the Board of Directors of Glencoe Software and a member of the Scientific Advisory Board for RareCyte and Montai Health; he holds equity in Glencoe and RareCyte.

M.P.S. is a cofounder and scientific advisor of Crosshair Therapeutics, Exposomics, Filtricine, Fodsel, iollo, InVu Health, January AI, Marble Therapeutics, Mirvie, Next Thought AI, Orange Street Ventures, Personalis, Protos Biologics, Qbio, RTHM, SensOmics. M.P.S. is a scientific advisor of Abbratech, Applied Cognition, Enovone, Jupiter Therapeutics, M3 Helium, Mitrix, Neuvivo, Onza, Sigil Biosciences, TranscribeGlass, WndrHLTH, Yuvan Research. M.P.S. is a co-founder of NiMo Therapeutics. M.P.S. is an investor and scientific advisor of R42 and Swaza. M.P.S. is an investor in Repair Biotechnologies.

The other authors declare no competing interests.

## Notes

https://cns-iu.github.io/hra-organ-gallery-supporting-information

https://cns-iu.github.io/hra-organ-gallery-supporting-information/#videos

https://github.com/cns-iu/hra-organ-gallery-supporting-information

https://github.com/cns-iu/hra-organ-gallery-in-vr-preprocessing

### Summary of Updates

In this revision, the length of the Methods section has been adjusted for a leaner reading experience. Code and other documentation has been moved to the Supplementary Information.

