## Supplementary Information for "Multiscale harmonization, semantic integration, and immersive exploration of single-cell data in support of novel biological insights"

### Supplementary Figures

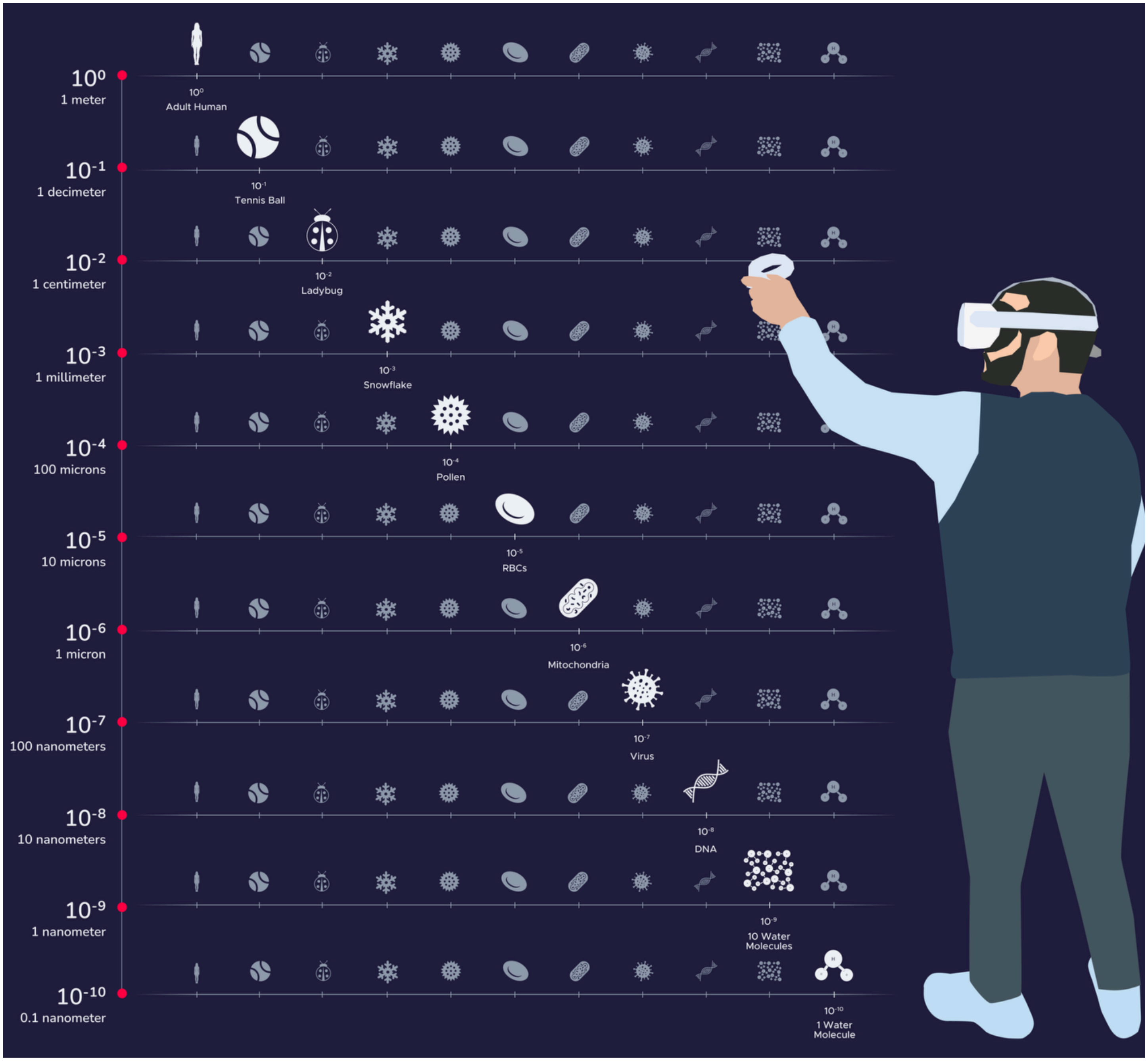


**Supplementary Fig. 1. Skyboxes for scenes on every level show a numbered line with the power ($10^x$) and a representative Icon for an object at that scale.** When laid out next to each other, they look like constellations in the night sky.

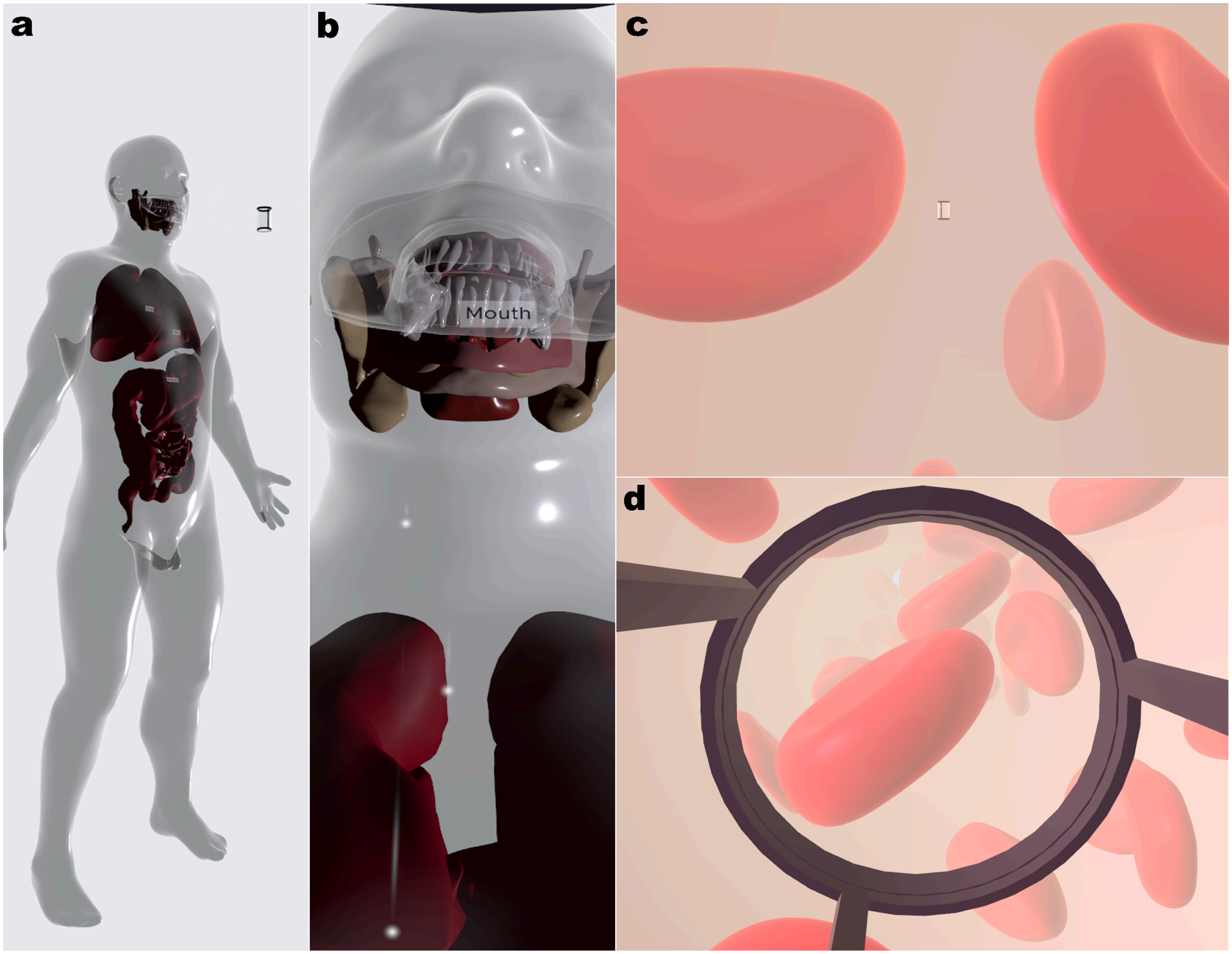


**Supplementary Fig. 2. The 2 reference objects for the Elevator Transition Scenes while traveling to the $10^{-2}$ (a-b) and $10^{-7}$ levels (c-d).**

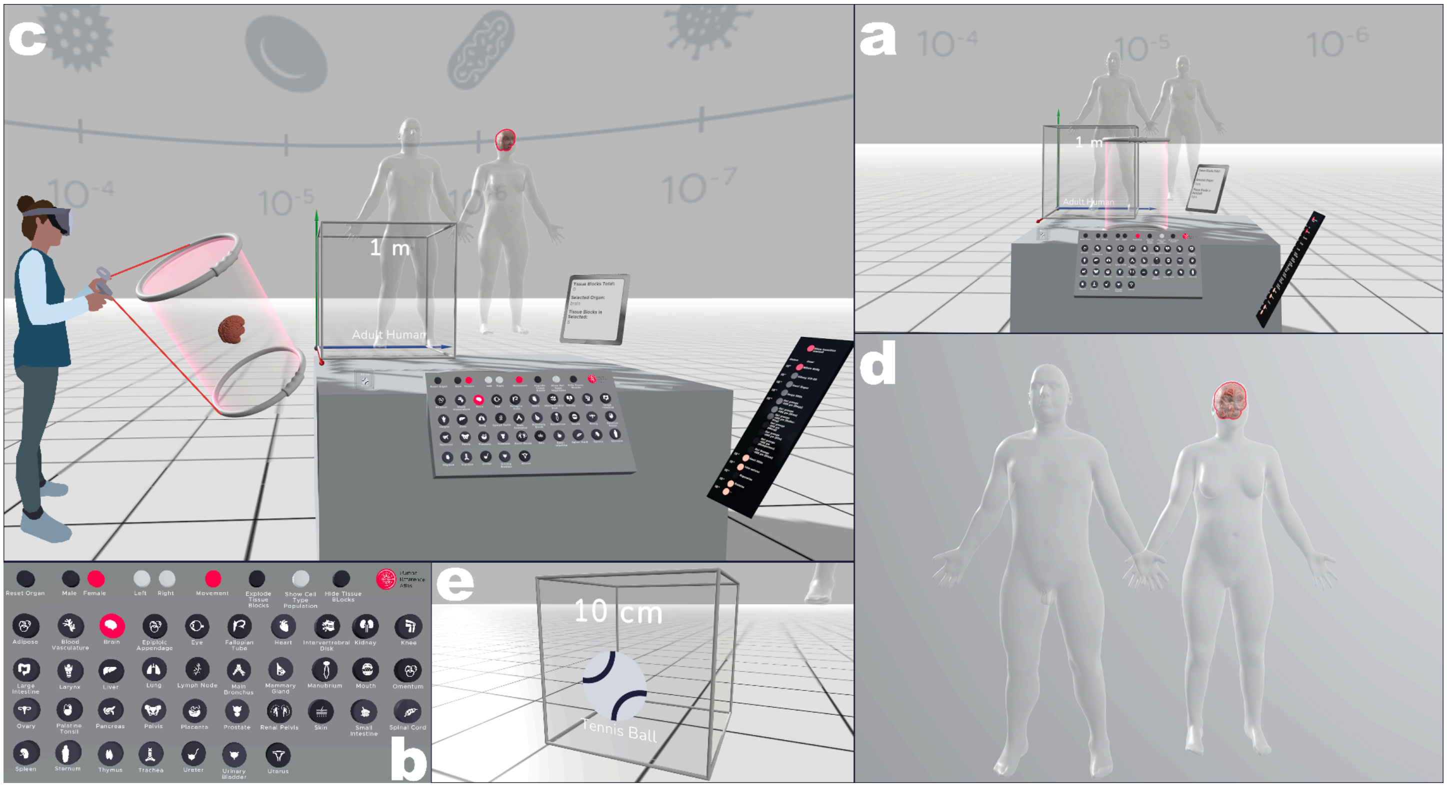


**Supplementary Fig. 3. a, Overview of hra-whole_body-hra-1_meter-10_0 with work area and Skybox overhead. b,** Organ selection keyboard with organ and sex selection buttons (brain, female) selected. **c,** The user holding the grab cylinder with the female brain. **d, The** currently selected organ is highlighted inside the male and female reference bodies. **e,** Detail shot of the small scale cube on the left side of the user.

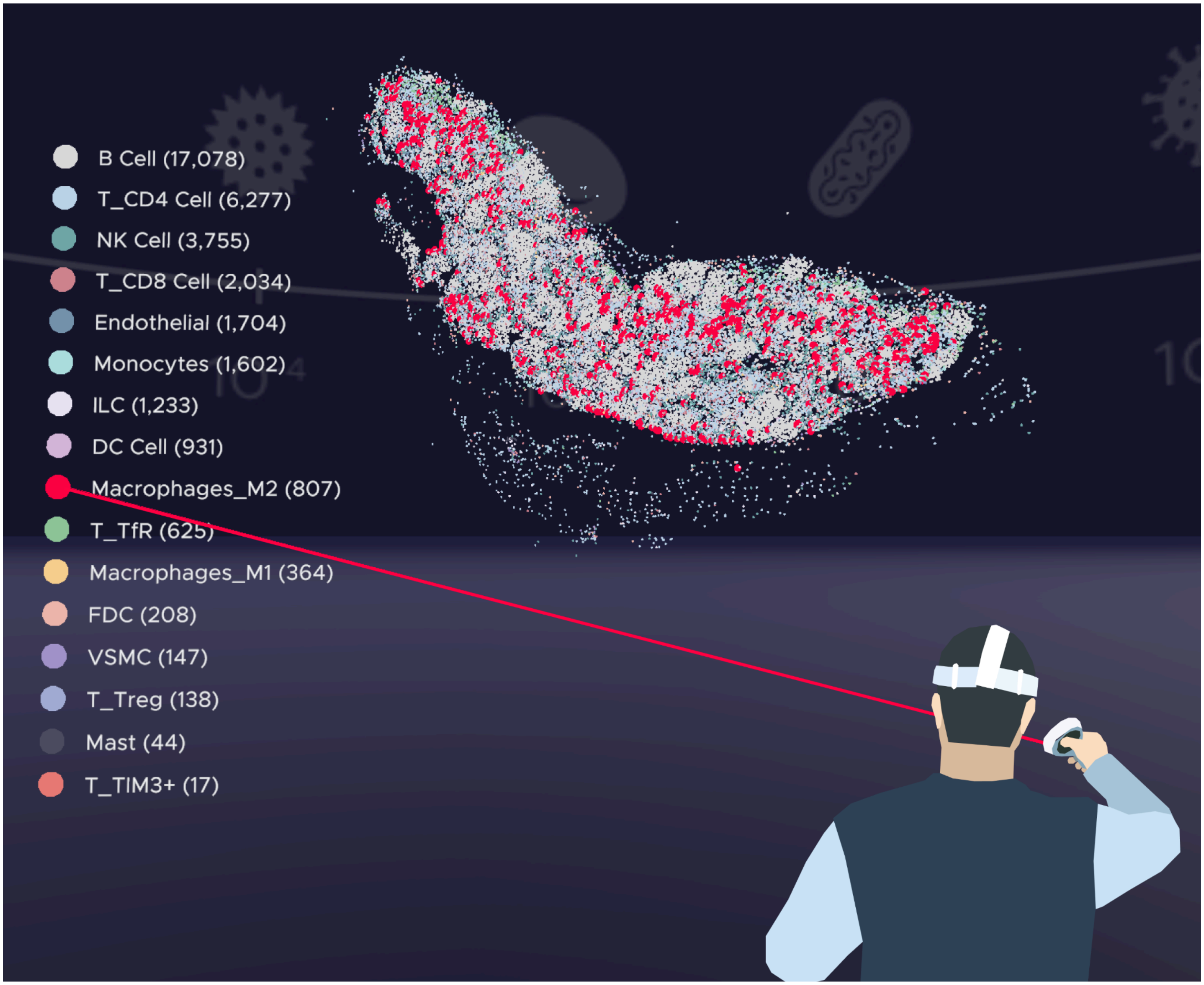


**Supplementary Fig. 4. A user interacting with the senescence-focused lymph node dataset in the sennet-lymph_node-enninful_farzad-1_centimeter-10_2 scene.** As they hover over the macrophages entry in the legend, the corresponding cells (in red) are highlighted (brush-and-link).

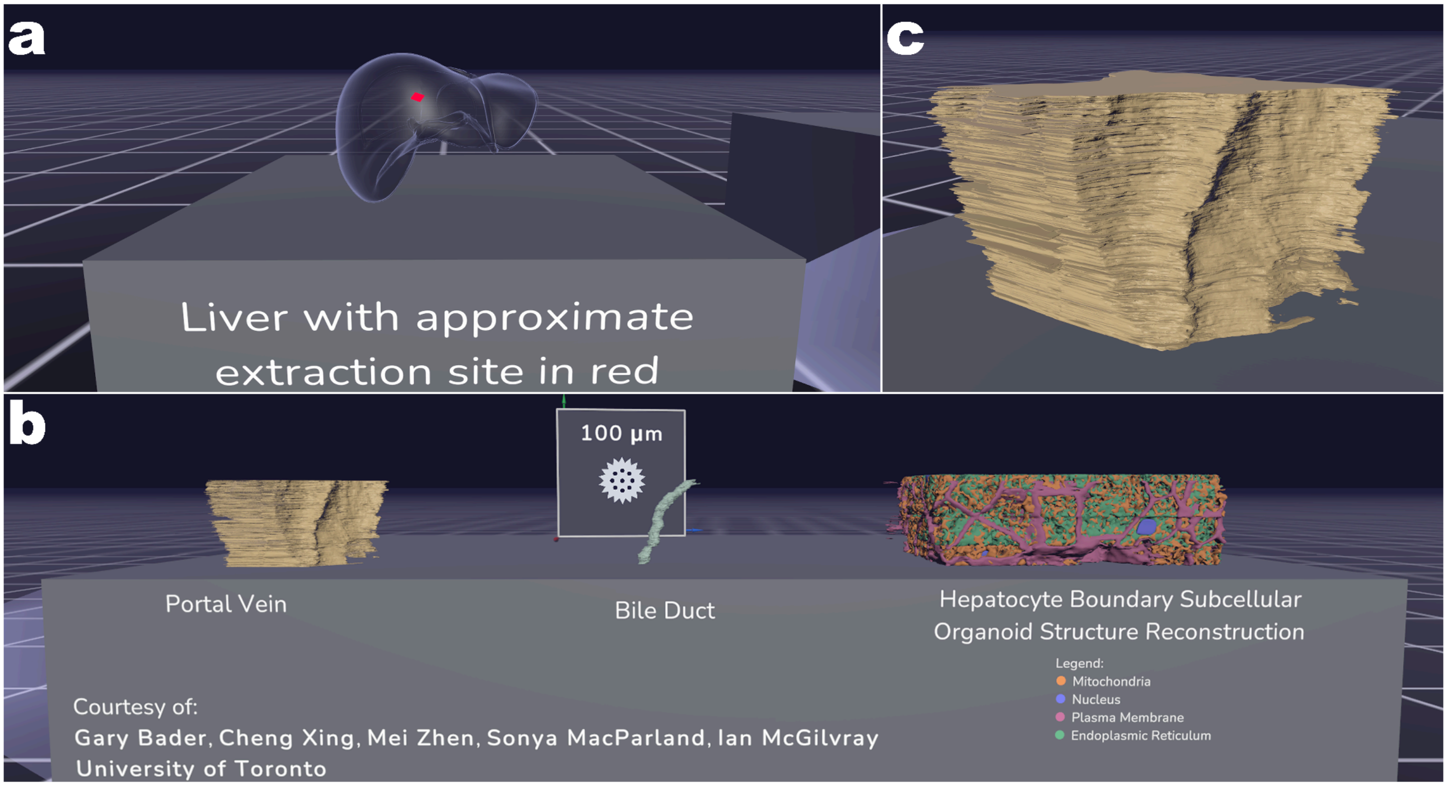


**Supplementary Fig. 5. Detailed views of the cifar-liver-bader_xing-100_microns-10_4 scene. a,** Extraction Site Component for the periportal region of the liver. **b,** Portal vein, bile duct, and hepatocyte organelle structures on a table next to each other, each of them grabbable, movable, rotatable, and scalable. **c,** Detail view of the portal vein. Future work aims to improve the sectioning artifacts visible on the surface of the model.

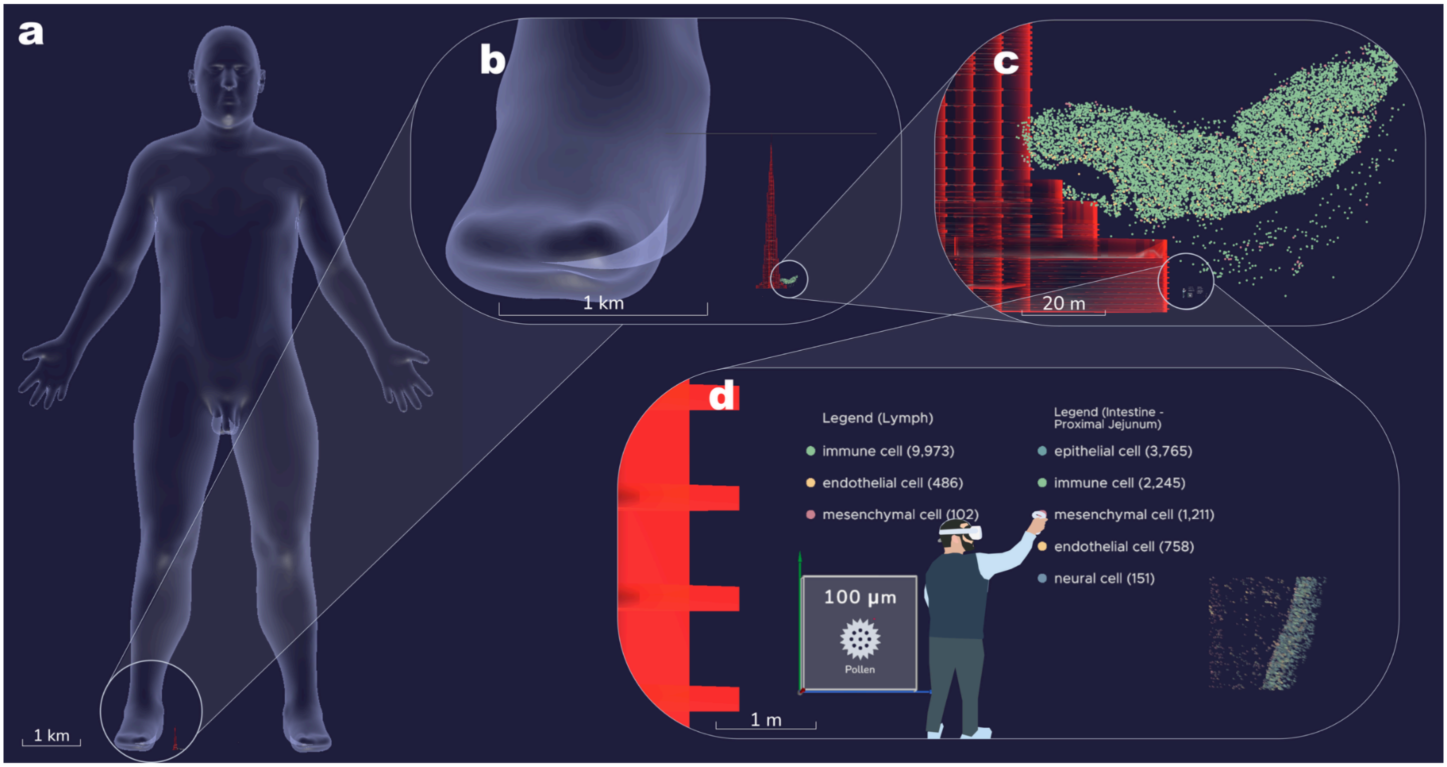


**Supplementary Fig. 6. Multiscale view of the datasets in hubmap-small_intestine-miao-100_microns-10_4 and sennet-lymph_node-enninful_farzad-1_centimeter-10_2 with orthographic rendering and scale bars to enable comparison in hra-multiscale-comparison-codex-100_microns-10_4**. **a,** The whole human is visible at four orders of magnitude larger than the user and the data, with Burj Khalifa (red) in its normally scaled height (828 meters) barely visible by the whole human's right foot. **b,** A zoom into the whole human's right foot with Burj Khalifa now visible. Note that the building's spire barely reaches the top of the whole human's foot. **c,** Zooming in further, the lymph node data now prominently appears at the base of Burj Khalifa. **d,** The user interacts with the shared legend while the small intestine data and 100 micron Scale Cube are fully visible (1x1 meter).

### Supplementary Note 1: Workshop

The collaborative development of the HRA: Powers of Ten module was facilitated through a HuBMAP JumpStart Fellowship (hubmapconsortium.org/jumpstart-program/#andreas2024), which enabled an in-person workshop, hosted by the Bioinformatics and Computational Biosciences Branch ( www.niaid.nih.gov/research/bioinformatics-computational-biosciences-branch) at the National Institute of Allergy and Infectious Diseases in October 2024. This allowed for dialogue and collaboration between early-stage investigators (ESIs) and BCBB experts in structural biology, medical illustration, and virtual reality (VR) regarding collaborative identification of user needs, immersive visualization, and 3D modeling. Initial ideas for interacting with single-cell data in VR across scales were implemented and have since been developed further, leading to the results documented in this paper.

#### Workshop website

Information about goals, attendees, organizers, and photos from the HRA: Powers of Ten Workshop is available at cns-iu.github.io/workshops/2024-10-24-jumpstart-workshop.

#### Workshop abstracts submitted by ESIs

*Alex Yu Hin Wong, Ph.D.*

**Abstract**: Advancements in data collection and computational methods are essential for fully describing and quantifying the intricate organization and biology of healthy and diseased tissues. Understanding key features such as immune-cell interactions, epithelial cell morphology and state, and macroscale structures like supporting vessels and nerves requires comprehensive analysis at multiple spatial scales. These complexities necessitate innovative approaches to data acquisition, processing, and visualization to unravel the underlying biological mechanisms and disease processes. In my current research, I am collecting and analyzing a fundamentally new type of high-plex three-dimensional (3D) data from normal and diseased colon tissues. By referencing both the Human Tumor Atlas Network (HTAN) and the Human BioMolecular Atlas Program (HuBMAP) ontologies, I aim to create a detailed map of cellular interactions and tissue architecture. The samples I work with are up to several millimeters thick and can be reliably imaged across many channels using advanced multiplexed light-sheet microscopy techniques. This allows for the capture of different spatial scales while maintaining single-cell resolution in each acquisition cycle. However, the sheer volume of data presents significant challenges. Each dataset, composed of multiple imaging cycles, can accumulate tens of terabytes. This massive data size makes it difficult to quickly inspect and analyze the information using traditional visualization and analysis tools. Rapid and efficient data inspection is crucial for identifying patterns, anomalies, and interactions that could lead to new insights into tissue biology and disease mechanisms. I am eager to attend the Powers of Ten workshop because it focuses on developing new virtual reality (VR) visualization methods and interfaces which will be essential for the comprehensive exploration of such large and complex multiplexed light-sheet datasets. The workshop provides a unique opportunity to collaborate with experts in VR visualization, data encoding, and biological data analysis. In particular, I am interested in exploring how data can be encoded into alternative representations, such as mesh models, which can significantly reduce storage space requirements while retaining all the necessary spatial information for analysis. My project specifically focuses on neuroimmune interactions within colon tissue. Understanding these interactions at both micro and macro scales is critical for elucidating the mechanisms of diseases such as colorectal cancer and inflammatory bowel disease. Furthermore, developing an “exploded view” mode in VR could greatly enhance our ability to highlight areas of interaction and inspect individual cells in detail. This functionality would allow researchers to deconstruct complex tissue architectures in a virtual environment, providing new perspectives and insights that are not possible with traditional two-dimensional (2D) imaging or even standard 3D visualization tools. From our experience with processing and visualizing 3D data, I recognize the potential of VR to revolutionize scientific visualization of multiscale datasets. By immersing researchers in a virtual environment, VR can facilitate a more intuitive understanding of spatial relationships and interactions within tissues. Collaborating with the Human Reference Atlas (HRA) VR working group at the workshop could help bridge the gap between data generation and meaningful interpretation, leading to the creation of an intuitive VR visualization platform for 3D spatial omics. Attending the Powers of Ten workshop will enable me to share my insights and challenges with a community of like-minded researchers and technologists. I hope to contribute my experience in high-resolution imaging and data processing to the development of innovative VR visualization tools. I also hope to learn from the expertise of others in the fields of VR interface design, data compression, and multiscale visualization techniques.

*Yang Miao*

**Abstract**: Working in a spatial proteomics lab, my research focuses on understanding the complex networks of cell interactions through multiplexed imaging techniques, such as CODEX. I am eager to attend the workshop to deepen my understanding of advanced biovisualization techniques and explore how these tools can be applied to the analysis of spatial-omics datasets. One of my key interests is learning how to effectively visualize the datasets in 3D. Integrating spatial data with current virtual reality (VR) technology offers a unique opportunity to explore organs and tissues at their corporal locations. It provides an interactive and user-friendly platform for analyzing intricate spatial datasets; I am particularly excited to learn how VR can be leveraged to uncover new biological insights. Understanding the current challenges in biovisualization is another area of focus for me. The hands-on sessions with VR devices are an exciting aspect of the workshop, as they will provide practical insights into how these technologies can be integrated into existing data analysis workflows. Incorporating these tools into regular research practices could significantly enhance the interpretation of spatial datasets, benefiting both researchers and clinicians alike. In addition, I look forward to sharing our expertise in visualizing the hierarchical organization of tissues, including cell types, tissue neighborhoods, and broader community structures during the workshop. Developing multiscale visualizations of these elements could yield novel perspectives on tissue architecture and spatial relationships, making the exploration of multiplexed imaging data more intuitive and impactful. Thus, I am especially interested in how to optimize the techniques to highlight multiscale cellular neighborhoods in 3D visualizations. A further goal is to connect with fellow Junior Investigators in the HuBMAP community, fostering a collaborative environment where ideas and feedback on visualization techniques can be exchanged. This interaction will help me better understand how others in the field are addressing similar challenges in spatial-omics research. By attending this workshop, I aim to gain insights into the latest advancements in 3D biovisualization, along with practical skills that can be directly applied to my work with HuBMAP data. In return, I hope to contribute our experiences with spatial multiplexed imaging and multiscale analysis while engaging in meaningful discussions about the future of data visualization in spatial omics.

*Archibald Enninful, Ph.D.*

**Abstract**: Advances in spatial multiomics have transformed our ability to study tissue architecture, capturing details of cellular interactions and molecular states. However, the vast amount of data generated—often terabytes per sample—poses significant challenges for visualization and analysis. Virtual Reality (VR) offers an immersive platform to explore spatial multiomic datasets. Techniques like CODEX and spatial transcriptomics capture the spatial distribution of proteins and RNA within tissues at single-cell resolution. In my research with the NIH's Cellular Senescence Network (SenNet) and the Human Tumor Atlas Network (HTAN), I focus on mapping the lymph node microenvironment and senescent cells. Senescence, a process linked to aging and cancer, requires spatial context to fully understand its impact on tissue behavior. Traditional 2D imaging often fails to capture these important dynamics, especially in complex tissues like lymph nodes and tumors. VR addresses this challenge by enabling 3D exploration of tissue architecture, providing new ways to visualize cell-cell interactions and tissue structure. By rendering features such as immune cells, epithelial structures, and vasculature, VR offers an intuitive platform for analysis. In my work, VR allows detailed examination of senescent cells within lymph nodes, revealing patterns that may be missed through 2D methods. Overall, VR is a powerful tool for visualizing spatial multiomic data, enabling immersive exploration of tissue structures and uncovering new insights into processes like senescence and cancer. I look forward to connecting with fellow Junior Investigators across NIH consortia to further explore the use of VR for spatial omics data visualization.

*Chenchen Zhu, Ph.D.*

**Abstract**: Mapping the precise architecture of human tissues in a preserved 3D environment is key in understanding normal tissue functions and how they are subverted in disease. Spatial transcriptomics provides deep insight into cellular states and programs and is increasingly capable of single-cell analysis, particularly when dissociative single-cell data is also available. Recently, we have developed an approach to reconstruct 3D tissue maps for the human gut from serial thin sections profiled with spatial transcriptomics or high-plex immune-based protein imaging. We have discovered that few if any cells in traditional 5 μm sections are complete, substantially complicating accurate phenotyping. Moreover, 2D images miss key features of tissue ultrastructure that are readily detected in 3D. These observations underscore the importance of 3D tissue mapping using highly multiplexed assays. Our approach using serial section reconstruction is the least "sophisticated" approach for assaying 3D molecular maps, but it has the great advantage of enabling multi-omic analysis, for example with one set of sections devoted to transcript profiling and an interleaved set to protein-based imaging. Some spatial transcriptomic methods are also compatible with subsequent protein-based tissue imaging. The key challenge is to reconstruct the 3D model from these serial images through image or feature based alignment. To this end, we have built a robust 3D-reconstruction method called Space-Map that aligns tissue sections using multimodal features and reconstructing detailed tissue maps. To appreciate the stereotyped organization of the normal colon, 3D visualization is crucial. My focus for the workshop is to collaborate with the team to find the best ways to visualize our 3D models at multiple levels—including tissues, cells, and molecules (RNA). These represent vastly different scales in biology and are essential for understanding tissue biology. Using 3D rendering in virtual reality will be key to delivering the stereoscopic sense of the models, helping us improve our 3D reconstruction approach. To assess the 3D model, I will specially focus on stem cells and their descendants in the colonic villus that are involved in the normal function of the intestinal mucosa. The mucosa is also rich in immune cells, some of which are distributed through the tissue and others of which are concentrated in secondary lymphoid structures in Peyer's patches. We will pay particular attention to the analysis of these diverse tissue-resident immune cells with the goal of more precise and accurate cell-type calling. In summary, this VR workshop will connect me with other 3D map builders working on various tissues within HuBMAP. By utilizing the power of VR, I aim to assess our 3D models and uncover new biological discoveries that are not readily visible with 2D rendering. Additionally, I am personally very interested in learning and understanding VR development and hope to apply this technology to other genomic visualizations.

### Supplementary Note 2: Software architecture of the HRA Organ Gallery

The HRA Organ Gallery is built with Unity 2022.3 and the XR Interaction Toolkit 2.6. The XR Interaction Toolkit handles all VR input: users point at objects and UI elements with a ray from their controllers, grab and move objects (*XRGrabInteractable*), scale them (*XRGeneralGrabTransformer*), and snap them back to their original position, rotation, and scale (*socket interactors*). 2D text, legends, and buttons are rendered on Unity Canvas elements placed in 3D space.

All data displayed in the application is converted during development into *ScriptableObjects*, i.e., serialized data containers stored as assets in the Unity project, so the application runs without an internet connection. The main *ScriptableObject* types are:

*SOCellPositionList* and *SODatasetCellTypeFrequency*: cell positions (x, y, z) with cell type labels, and the number of cells per type, ingested from CSV files.

*SOCellColorMapping*: mappings from cell type labels to colors, shared across scenes.

*SOLevelIndex* and *SOLevelList*: scene metadata (name, scale, Skybox, Icon) that the Elevator Panel uses to load scenes.

*SONodeArrayFromAPI*: cached HRA API responses with 3D reference organs and tissue blocks, updated with each HRA release.

Developer documentation is available at [cns-iu.github.io/hra-organ-gallery-supporting-information/#native-unity-packages-and-concepts](cns-iu.github.io/hra-organ-gallery-supporting-information/#native-unity-packages-and-concepts).

Custom components and Editor tools that build on these types are described in **Supplementary Note 3**.

### Supplementary Note 3: Reusable components and Editor tools

On top of the Unity framework described in **Supplementary Note 2**, the HRA Organ Gallery uses a small set of custom C# components that are shared across scenes, and 2 Editor tools that prepare data before the application is built.

**Visualization**: *VisualizerBase* is an abstract base class that places a visualization inside a container in 3D space with a given maximum width, scaling factor, and color scheme. Two derived classes implement it. *Visualizer3D* renders cells as colored dots from a list of positions and cell types in 2D or 3D; it is used in the lymph node, small intestine, large intestine (zhu), and multiscale comparison scenes. *VisualizerBiomarkers* renders Visium spots on a floor-aligned plane with vertical lines for senescence hallmark expression; it is used in the brain scene. Each dot is an instance of a shared prefab carrying a *CellData* component that stores the cell's label and color, so they can be queried at runtime.

**Legends and brushing and linking**: *CellLegend* and *LegendDisplay* build a 3D legend from the cell type frequencies and color mapping of a dataset. When the user points at a legend entry with a VR controller, OnHoverSendMessage broadcasts that cell type label and all matching cells in the scene are highlighted.

**Editor tools**: *IngestCellPositions* converts CSV files of cell positions and types into *ScriptableObjects*, which load faster than reading CSV at runtime. *IngestScene* caches 3D reference organs and tissue blocks from the HRA API as *ScriptableObjects*; it is run before each HRA release (June and December) so the application works offline.

Developer documentation is available at [cns-iu.github.io/hra-organ-gallery-supporting-information/#custom-c-components-and-prefabs](cns-iu.github.io/hra-organ-gallery-supporting-information/#custom-c-components-and-prefabs) and [cns-iu.github.io/hra-organ-gallery-supporting-information/#editor-scripts-and-editorwindows](cns-iu.github.io/hra-organ-gallery-supporting-information/#editor-scripts-and-editorwindows).

### Supplementary Note 4: Matching 3D cell meshes to the quantification table in hubmap-large_intestine-wong-100_microns-10_4

For the light-sheet crop (**Fig. 6.a**, left), co-author Wong provided a quantification table (CSV) and a 3D model of CD8 cells (OBJ). The quantification table had 1 row per cell with an ID, position, and volume, whereas the 3D model used generic object names (e.g., "CD8.001", "CD8.002") that did not match the table IDs. Manual mapping was impractical because of the number of cells and because many were located deep inside the 3D structure, which made them hard to identify by eye. A Python script was therefore written to match each 3D object to its row in the table using both position and volume.

**Reference cell and coordinate offset**: To map the table's coordinate system onto that of the 3D model, 1 easily identifiable cell (cell_086, renamed manually beforehand) was used as a reference point. The coordinate offset was calculated by subtracting the reference cell's position in the table from its position in the 3D model:

$$\Delta_p = p_{3D} - p_{CSV}$$

**Unit conversion**: Table coordinates were in microns, but the units of the 3D model were unknown. The script therefore tested 3 conversion factors: microns to meters (1e-6), microns to millimeters (1e-3), and no conversion (1.0). It selected the factor that produced the most successful matches.

**Matching**: For each cell in the 3D model, the script (1) converted the CSV position using the selected unit conversion and offset, (2) calculated the distance d between the 3D cell and the converted CSV position, and (3) accepted a match only if both position and volume criteria were met:

$$Match = 1,\ if\ d \le d_{thresh}\ and\ \Delta V \le \Delta V_{thresh};\ otherwise\ 0$$

where

$$\Delta V = \left|\frac{V_{Blender} - V_{CSV}}{V_{CSV}}\right| \times 100$$

Mesh volumes (*V_Blender*), measured in cubic scene units, were computed with *bmesh* (docs.blender.org/api/current/bmesh.html), a built-in Blender library, accounting for object scaling and transformations. The thresholds were set by trial and error to d_thresh = 3.0 scene units and ΔV_thresh = 20%. Once a CSV ID was matched, it was removed from the pool of candidate IDs so no two 3D cells could claim the same ID. Match quality metrics were logged.

**Validation**: The Data Provider (co-author Wong) validated the resulting mapping. For example, 3D object CD8.001 was matched to cell_023 with d = 1.24 scene units and ΔV = 11.7% (V_Blender = 0.001234, V_CSV = 0.001398), both within their thresholds.

### Supplementary Note 5: Scene implementation details for hubmap-large_intestine-wong-100_microns-10_4

**Scene setup:** After cell matching (**Supplementary Note 4**), the 3D model with renamed cells was exported as FBX and imported into Unity. Light probes placed around the model provide mixed lighting to balance visual quality and VR performance. Users can fly freely in any direction to orbit the cell structure, pass through dense regions, and move inside the tissue volume (**Fig. 2.b**).

**Cell sockets and tethering:** Each cell has a socket interactor (**Supplementary Note 2**) at its original position; cells released within 0.3 meters of it snap back into place. A custom socket accepts only the cell whose ID matches its own, so cells cannot be returned to the wrong location. While a cell is grabbed, a white tether line connects it to its original position (**Fig. 6.b**) and disappears when the cell is returned.

**Heatmap and info panel:** A dropdown colors cells from yellow (low) to red (high) by minimum intensity, volume, distance from origin, or sphericity (**Fig. 6.c-d**), using values from the quantification table stored as a *ScriptableObject* (**Supplementary Note 2**). When a cell is grabbed, a panel shows its position (x, y, z), minimum intensity, volume, and sphericity.
